# Validating genuine changes in Heartbeat Evoked Potentials using Pseudotrials and Surrogate Procedures

**DOI:** 10.1101/2024.08.24.609348

**Authors:** Paul Steinfath, Nadine Herzog, Antonin Fourcade, Christian Sander, Vadim Nikulin, Arno Villringer

## Abstract

The brain continuously receives interoceptive information about the state and function of our internal organs. For instance, each time the heart beats, the brain responds by generating time-locked activity, known as heartbeat evoked potentials (HEP). When investigating HEPs, it is essential to adequately control for heartbeat-independent confounding activity to avoid false interpretation. In the present study, we highlight the pitfalls of uncontrolled analyses and advocate for the use of surrogate heartbeat analysis and pseudotrial correction, which are promising tools to control for spurious results. Surrogate heartbeat analysis involves shuffling the timing of heartbeats to verify the time-locking of HEP effects. Pseudotrial correction works by subtracting heartbeat-independent activity from HEPs. In this study we employ both procedures, validate them in simulations and apply them to real EEG data. Using EEG recordings obtained during the performance of an auditory novelty oddball task in a large population, we show that, without control analyses, pre-stimulus HEPs appear inversely related to task-related measures such as P300 event-related potential amplitudes and reaction time speed. However, these effects disappear after carefully controlling for heartbeat-unrelated EEG activity. Additionally, in real and simulated data, we show that pseudotrial correction has the potential to remove task-related confounds from HEPs, thereby uncovering real heartbeat-related effects that otherwise could be missed. This study therefore highlights issues that can arise when analyzing HEPs during tasks, provides solutions to overcome them, and gives recommendations for future studies to avoid pitfalls when analyzing and designing behavioral paradigms with HEPs.

## Introduction

Interoception, the sensing of internal bodily states, is a fundamental aspect of physiological and psychological functioning. The brain constantly monitors and adjusts the state and function of our internal organs, including hunger, thirst, respiratory, and cardiac functions (Craig, 2002). Research has shown that the neural processing of visceral signals contributes to a wide range of emotional, perceptual, and cognitive processes (Azzalini, Buot, et al., 2019; Critchley & Harrison, 2013; Quigley et al., 2021). The heart, in particular, is a crucial source of interoceptive information. Importantly, neural responses to heartbeats can be directly measured by neuroimaging modalities such as EEG or MEG. These heart-evoked potentials (HEP) are event-related potentials (ERP) time-locked to participants’ heartbeats (Schandry et al., 1986).

The HEP appears to reflect the cortical processing of afferent signals generated by the heart and is commonly used as an objective neurophysiological marker for cardiac interoception (Coll et al., 2020; Jones et al., 1986; Pollatos & Schandry, 2004; Schandry et al., 1986). Various studies have demonstrated that HEP amplitudes are significantly higher during interoceptive compared to exteroceptive attention (Montoya et al., 1993; Petzschner et al., 2019; Pollatos et al., 2005; Pollatos & Schandry, 2004; Schandry et al., 1986; Villena-González et al., 2017), suggesting that increases in its amplitude indicate an attentional shift towards internal stimuli (Al et al., 2020). While this relationship depends on the individual ability to feel the heartbeat (Yuan et al., 2007), it can be further strengthened with training of cardiac perception (Canales-Johnson et al., 2015). Even though heartbeats are often not explicitly consciously processed, the brain nonetheless continuously generates HEPs, indicating often unconscious processing of heartbeat-related information (Kern et al., 2013; Perogamvros et al., 2019).

Importantly, when HEPs are investigated during tasks, controlling for the concurrent task-evoked activities is essential to ensure that the measured HEPs accurately reflect heartbeat evoked responses. If HEPs overlap with task-related evoked activity, differences in HEPs can occur solely due to differences in evoked activity. In this context, a surrogate heartbeat control analysis has been suggested (Azzalini et al., 2019; Babo-Rebelo et al., 2016; Park et al., 2014; Park & Blanke, 2019; Park et al., 2016). This control analysis operates on the premise that effects genuinely related to the heartbeat should be time-locked to it. To test for this time-locking, surrogate heartbeats are generated by perturbing the original timing of the HEP onsets (i.e. by randomly shifting the R-/T-peak markers in time (Park et al., 2016) or exchanging the heartbeat onset timing across trials Azzalini et al., 2021). This procedure is repeated many times, each time performing the same statistical analyses with the surrogate datasets. The results are then used to generate a null-distribution to which the original HEP effects are compared. The main idea is that the heartbeat onset shuffling should preserve all other aspects of neuronal data not directly related to HEPs. If the original effect exceeds the effects obtained by analyzing surrogates, it likely reflects a genuine heartbeat-related process (Park et al., 2014). However, a theoretical limitation of the surrogate heartbeat analysis is that true HEP effects may remain undetected if they are mixed with confounds (type II error). In order to overcome this challenge, in this study, we thus combine a surrogate control analysis with a pseudotrial correction (Wainio-Theberge et al., 2021) and show the potential to uncover genuine HEP effects. The pseudotrial correction procedure involves adding random triggers to the data with no time-locking to the heart, which allows to estimate activity unrelated to the heartbeat (e.g. task-related responses). This results in pseudotrials which can be subtracted from real HEPs, ideally removing confounds while revealing true HEP effects.

In the present study, we demonstrate that the surrogate heartbeat analysis as well as pseudotrial correction can effectively control for confounding factors. Using a large population-based EEG dataset recorded during an auditory novelty oddball task (Loeffler et al., 2015) we observed a significant inverse relationship between the amplitudes of pre-stimulus HEPs and task-evoked P300 ERP - an ERP generated in response to the target stimulus which is assumed to reflect allocation of attentional resources (Kok, 2001; Polich, 2007). Furthermore, we observed lower HEP amplitudes before trials with faster reaction times. While these observations are in line with the idea that lower HEP amplitudes reflect the allocation of attentional resources towards exteroception (Al et al., 2021; Al, et al., 2020; Marshall et al., 2019, Zaccaro et al., 2022), the application of surrogate heartbeat and pseudotrial correction showed that these effects are rather spurious in our task. By utilizing resting-state as well as simulated data, we present alternative explanations not involving HEP related effects. Furthermore, using real and simulated data, we show that pseudotrial correction can uncover HEP effects which otherwise are masked by confounding factors, thereby reducing the chance of false negative findings. Our study underscores the need to apply careful control analyses when investigating the relationship between cardiac- and task-evoked activity and represents a significant advancement towards the robust examination of HEP effects. The suggested procedures provide an avenue for a more comprehensive understanding of the neurophysiological mechanisms through which HEPs may influence cognitive processes, and encourages a critical review of previous findings.

## 2. Methods

### 2.1. Participants

The data used in this study were recorded as part of the population-based LIFE-Adult study (Leipzig Research Center for Civilization Diseases, Leipzig University; Loeffler et al., 2015). Participants were chosen at random from the residence registration office, and written informed consent was from the participants. The participants received monetary remuneration. The study was approved by the University of Leipzig’s Medical Faculty’s ethics committee. EEG data was available from 2887 subjects. Only subjects with task and resting state recordings were included. We excluded subjects with a history of brain hemorrhage, concussion, skull fracture, brain surgery, or brain tumor. Only right-handed people were included. No alcohol consumption was allowed on the day of the measurement. Participants who consumed central nervous system affecting medications were excluded (opioids, hypnotics and sedatives, anti-parkinsonian drugs, anxiolytics, anti-depressants, anti-psychotics, anti-epileptic drugs).

In addition, subjects with less than 20 heartbeats in the pre-stimulus time of the target stimuli were excluded (see EEG data processing, Heartbeat Evoked Potentials). After applying these criteria, our final sample included 1739 participants’ datasets (mean age = 70, SD = 4.6, 874 females).

### 2.2. EEG recording

EEG data was recorded from 31-channel Ag/AgCl scalp electrodes (Brain Products GmbH, Gilching, Germany) in an electrically shielded and soundproof EEG booth. The electrodes were mounted in an elastic cap (easyCAP, Herrsching, Germany) according to the international standard 10–20 extended localization system. The signal was amplified with a QuickAmp amplifier (Brain Products GmbH, Gilching, Germany). Additionally, two electrodes recorded vertical (vEOG) and horizontal (hEOG) eye movements above and beneath the right eye. One bipolar electrode attached to the right and left forearm recorded electrocardiogram (ECG). The electrodes were referenced to the common average reference, with AFz being a ground electrode. The electrodes’ impedances were kept below 10 kΩ, the sampling rate was 1000 Hz, and the data was low-pass filtered at 280 Hz. At the start of the EEG measurement, the participants underwent a 20-minute-long resting state EEG (rsEEG) measurement. During this time, they laid on their back in a dark room and closed their eyes. The participants were instructed to remain awake and avoid falling asleep. This was followed by a 15-min-long auditory novelty oddball paradigm described below (2.4). A more detailed description regarding the recording can be found in a paper by Jawinski and colleagues (Jawinski et al., 2017).

### 2.3. EEG data processing

EEG processing was performed with MATLAB (version R2024a) using custom-written scripts as well as the EEGLAB (Delorme & Makeig, 2004), Fieldtrip (Oostenveld et al., 2011), HEPLAB (Perakakis, 2019), and ERPLAB (Lopez-Calderon & Luck, 2014) toolboxes. EEG data were filtered with a 4th order Butterworth filter, first high-pass with a 1 Hz cutoff and subsequently low-pass with a 45 Hz filter cutoff frequency. In addition, to reduce residual line noise, a notch filter between 49 and 51 Hz was applied using the EEGLAB pop_eegfiltnew function. The data was down-sampled to 250 Hz. Bad channels were identified and removed using the clean_artifacts EEGLAB plugin if they were flat for more than 5 seconds, correlated with their neighbors less than 0.85, or had residual line noise (average number of removed channels 0.94, *SD* = 1.15). Removed channels were interpolated using spherical interpolation. To identify noise sources in the data, we performed Independent Component Analysis (ICA) using the extended infomax algorithm after Principal Component Analysis dimensions reduction to the rank of the data. R-peaks were identified in the ECG data using the heplab_slowdetect function. ECG and independent component time-courses were segmented between -0.05 to 0.6 s around the R-peak markers, averaged across epochs, and correlated with each other. IC’s with a correlation greater than 0.8 with the ECG were marked as artifact (Al, et al., 2020). IClabel was used to identify potential components related to eyes (prob. ≥ 0.6), muscles (prob ≥ 0.5), line noise (prob. ≥ 0.5), channel noise (prob. ≥ 0.4) and other (prob. ≥ 0.5). The automatically marked components were reviewed manually, and the selection adapted if necessary. An average of 1 heart related ICs (*SD* = 1.08, min 0, max 5) were removed from the data.

The same preprocessing steps up until the ICA decomposition were repeated on the same data, but with a high-pass filter cutoff of 0.3 Hz. The ICA weights were copied to this data, and components marked as artifacts were removed. This was done, since ICA performs better with a higher high-pass filter cutoff, while slow ERP components could be degraded at such.

### 2.4. Oddball Task

A novelty auditory oddball paradigm was used to elicit auditory event related potentials. In total, 600 stimuli were presented in a pseudo-randomized order with a minimum of two standard stimuli between target stimuli, a maximum of nine standard stimuli in succession and a constant inter-stimulus interval (ISI) of 1500 milliseconds (ms). The more frequent, non-relevant standard stimuli (500 Hz sinusoidal tone lasting 40 ms including 10 ms rise and fall times) occurred 456 times with a global probability of 76%. Both the task-relevant target (1000 Hz sinusoidal tone lasting 40 ms including 10 ms rise and fall times) and novelty stimuli (environment or animal sounds lasting 400 ms with variable rise and fall times) occurred 72 times with a global probability of 12 % each. The participants were instructed to press a button when the target stimulus occurred. After 300 stimuli, the paradigm was interrupted by a short pause (30 s), in which the participant was asked to change the response hand. For the current study, we only analyzed target stimuli and their preceding HEPs. The participants performed the task with an accuracy of 96.18% correct trials, 3.65% incorrect responses, 0.16% omission errors, and an average reaction time of 489.6 ms.

**Figure 1.**
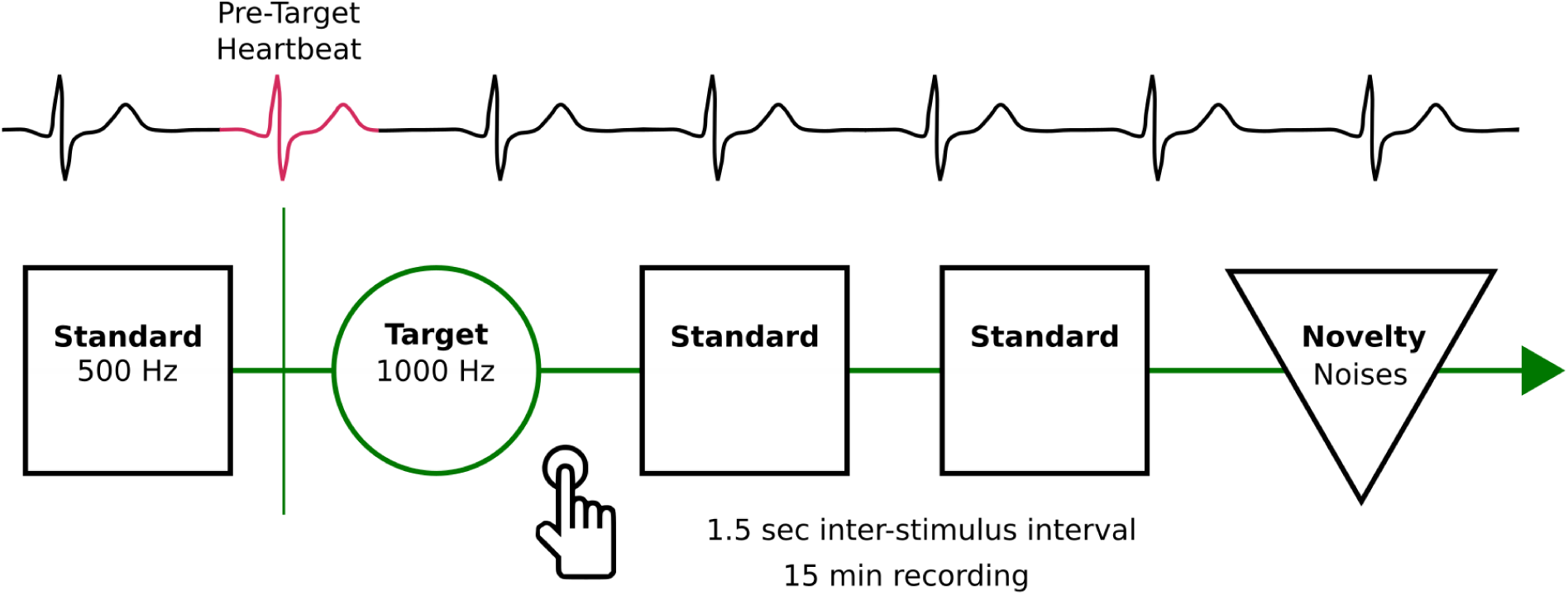
Schematic of the Auditory Novelty Oddball Task. Participants performed a 15-min-long auditory Novelty Oddball task in which they were presented with standard (500 Hz), target (1000 Hz), and Novelty (environment or animal) sounds. They were instructed to press a button each time the target stimulus occurred. Heartbeats in the pre-stimulus time of target trials were identified and used as marker for time-locked pre-stimulus heartbeat evoked potentials.

### 2.5. Heartbeat evoked potentials

We only selected pre-stimulus HEPs that occurred within the time window of -1100 and -600 ms before the target onset, in order to prevent interference from the preceding stimulus or the evoked response related to target stimulus presentation. HEPs were epoched between -200 and 600 ms around the R-peak. Epochs that exceeded a peak-to-peak signal amplitude ≥ 150 µV in any channel using moving window (window size = 200 ms, window step = 100 ms) were discarded. If fewer than 20 pre-stimulus HEPs were found, the subject was excluded from further analysis. As a result, on average, 27 (*SD* = 4.5) HEPs were included per subject. Baseline correction was applied based on the -150 to -50 ms time window. Due to the repetitive nature of the heartbeat, there is no ideal baseline window that is free of all activity (Park et al. 2014; Babo-Rebelo et al. 2016, 2019; Azzalini et al. 2019, Banellis and Damian 2020, Verdonk et al. in 2020) Hence, we repeated the same analyses without baseline correction and included the results in the supplementary materials.

### 2.6. ERP Analysis

Target trials were epoched between -500 to 1000 ms around the stimulus onset. Epochs that exceeded a maximum moving window peak-to-peak signal amplitude >= 150 µV were removed (window size = 200ms, window step = 100ms). To enhance the single-trial signal-to-noise ratio, we utilized a spatial filter derived from linear discriminant analysis (LDA) to distinguish between target and standard stimuli (Blankertz et al., 2010). For each participant, the average amplitude within the time window from 200 to 700 ms was calculated for target and standard ERPs, which yielded two amplitude values per participant. Using the amplitude values as features, we trained an LDA model to classify between the target and standard ERPs. The LDA model learns to maximize the difference between the two classes while minimizing the variance within each class. As a result, we obtain weights from the LDA model which reflect a spatial filter, which can be applied to the single trial data of each participant, resulting in a single P300 ERP time course per trial. To investigate how pre-stimulus HEPs relate to cognitive functioning, we sorted pre-stimulus HEPs into a) high vs. low P300 trials based on a median split of single trial average ERP amplitude within the 250–600 ms time window and b) in fast and slow category based on a median split of the subjects’ reaction times (RT).

### 2.8. Statistical analyses

We performed cluster-based permutation *t-*tests (two-tailed) on the data using the FieldTrip toolbox (Oostenveld et al., 2011) to compare HEPs between our conditions. The cluster-based permutation procedure is a method used to identify significant effects in time and space while accounting for multiple comparisons. This method works by adding up the *t*-values of all significant tests (*p*-value of < 0.05) that are adjacent both in space and time. These experimentally observed cluster-specific *t*-values are compared with a permutation distribution that is generated by randomizing the condition labels (in our case 1000 times) each time selecting the maximum summed *t*-value across all clusters. Clusters with a *p*-value smaller than an adjusted alpha level of 0.05 were considered significant. The precision of *p*-values derived from permutation tests is limited by the number of permutations. Since we applied 1000 permutations, the minimum *p*-value is 0.001. As previous studies are diverse in their temporal and spatial HEP findings, we included all 31 channels and the time window of -200 to 600 ms around the R-peak in our analyses. To control for volume conducted ECG artifacts, we compared the ECG signal related to the different conditions by temporal cluster-based permutation *t*-tests.

### 2.9. Surrogate heartbeat control analysis

To ensure that the observed HEP effects are genuinely linked to heartbeats and not a result of changes in ongoing spontaneous activity, we conducted a surrogate heartbeat analysis (Fig. 9A, Azzalini, et al., 2019). As a first step, we shuffled the pre-stimulus R-peak onsets within high & low P300 (fast & slow RT) condition. Then we performed the same statistical analysis using cluster-based permutations, while saving the largest summed *t*-value from the biggest identified cluster in positive and negative direction. Since in the surrogate data the temporal relationship between R-peak and HEP was broken by permuting the R-peak timings, the identified clusters should reflect heartbeat-unrelated processes. Since R-peak latencies were shuffled within subject and across conditions, the mean heart-rate and inter-beat interval is kept the same. We repeated this procedure 100 times, each time shuffling the pre-stimulus R-peak timings. At the end, we compared the cluster sum(*t*)-values from the original data with the corresponding distribution of the surrogate statistics. Since we test for positive and negative effects (two-tailed) the original *t*-value has to be larger than 97.5% or smaller than 2.5% of the surrogate cluster sum(*t*)-values to conclude a significant effect.

### 2.10. Pseudotrial correction

If genuine HEP effects are mixed with artifacts, the HEP effects can be missed because they are confounded by the artifact (see results of data simulation part 2.12). To uncover potentially present genuine HEP effects, we adopted a pseudotrial correction method suggested earlier for other purposes (Fig. 9B, Huang et al., 2017; Wainio-Theberge et al., 2021), which removes heartbeat independent effects. In this method, activity that is present in the pre-stimulus time of trials independent of a heartbeat is subtracted from real HEPs per condition, thereby effectively controlling for sorting induced pre-stimulus differences and regression to the mean effects (Bland & Altman, 1994). We implemented the pseudotrial correction in the following way: for each target trial a pseudo-R-peak-trigger was inserted at a random time point in the pre-stimulus time of interest (-1100 to -600 ms). Second, the data was sorted according to high & low P300 amplitude or fast & slow RT and averaged within these categories, resulting in pseudotrials. Lastly, within each subject, the real HEP epochs are corrected by subtracting the pseudotrials per condition.

### 2.11. Equivalence Test

For non-significant results, we employed equivalence testing to assess if the difference between conditions was smaller than a practically meaningful effect (smallest effect size of interest, SESOI). In contrast to classical null hypothesis testing, where the hypothesis is tested that a difference between groups exists, in equivalence testing the hypothesis is tested that the group difference is smaller than a SESOI. Hence, it is critical to define a reasonable SESOI which serves as the bound for effects to be considered too small to be meaningful (Lakens et al., 2018). Our HEP analyses are based on the idea that their amplitudes are modulated based on spontaneous fluctuations of interoceptive vs. exteroceptive attention. Hence, we chose to inform our SESOI by meta-analytic results from Coll et al. (2021) who investigated the relationship between HEP amplitudes and interoception across various publications in different domains (attention, performance, clinical, and arousal). From all reported effects across the different domains, we use the lowest end of the lowest confidence interval, which represents a Hedge’s *g* = 0.19 (performance category). Hence, we defined our SESOI as an effect smaller than a Hedge’s *g* of 0.19 and larger than -0.19. We used the two one-sided tests (TOST) procedure implemented in the R package TOSTER (version 0.8.0, *Lakens, 2017*).

### 2.12. Data simulation

To illustrate the increased power to detect effects when a pseudotrial correction is applied, we simulated data containing interactions between simulated HEP (simHEP) and task ERP (simERP). As a background signal with characteristic close to real data, we used EEG data of 100 random subjects from the Pz electrode which was scrambled by phase randomization to remove genuine activity (Gias, Carlos, 2024; Theiler et al., 1992). Phase randomization was achieved by first calculating the fast Fourier transform (FFT) of the original time-series signal, randomizing the phases, and performing the inverse FFT to reconstruct time-series data. As a result, genuine evoked responses (ERP and HEP) are removed from the signal, while oscillations at all frequency ranges remain intact.

Task triggers were added to this scrambled data every 4 seconds. We then simulated both positive and negative relationships between simHEP and simERP amplitudes with different magnitudes. This was achieved by first generating realistic normally distributed simERP amplitudes (mean = 7 µV, *SD* = 2) and simHEP amplitudes (mean= 1.5 µV, *SD* = 0.8). The signal-to-noise ratio (SNR, ratio of mean power between simulated evoked responses to the power of the phase randomized background signal) was 3.3 dB for simERPs and -9.3 dB for simHEPs. A scaling factor (k) between 0 and 1 for positive or 0 and -1 for negative relationships was used to represent the relationship strength between the two variables. To obtain simHEPs which are related to the simERP, we used the following approach: the normalized simERP amplitude was multiplied by the scaling factor adding the product of the square root of (1 - k²) and normalized simHEP amplitude. The resulting simHEP amplitudes were rescaled by their standard deviation and mean.

After generating correlated simERPs and simHEPs, they were modeled as positive deflections based on Gaussian functions in a single channel. The simERP onset was at 300 ms post-stimulus (duration 600 ms) and for half of the trials a pre-stimulus simHEP starting at 200 ms post stimulus (duration of 400 ms) was added at a random time between -1500 to -600 ms relative to the stimulus onset. Pseudotrials were defined as random triggers in the same pre-stimulus time of trials without simHEPs. On average, 421 trials (*SD* = 45) were present per simulated dataset.

We sorted the simERP trials based on their amplitude in the 250-600 ms time range, and tested for significant differences between the pre-stimulus HEPs using cluster-based permutation testing in the temporal domain. We then tested if a significant difference can be found after simHEPs were corrected by pseudotrial subtraction. A surrogate heartbeat control analysis was performed by randomly exchanging the onsets of pre-stimulus triggers per high & low amplitude condition before performing a cluster-based permutation test.

To adapt the surrogate procedure for pseudotrial corrected data, pre-stimulus triggers were randomly exchanged twice per high & low amplitude condition generating two sets of pseudotrials per condition. Next, the pseudotrials were averaged and subtracted from each other per condition, while the remaining activity was compared using cluster-based permutation testing. In this way, we control for insufficiently removed artifacts or potentially introduced noise by the pseudotrial correction procedure itself. This procedure was performed for each of the relationship directions (positive and negative association between pre-stimulus HEP and P300 ERP) and strength, and repeated 500 times to get a distribution of observed effects (maximum cluster-sum(*t*)) in each of the conditions.

Generate normally distributed simERP and simHEP amplitudes:

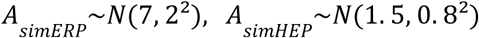

AsimERP and AsimHEP are the simulated ERP and HEP amplitudes. The notation *N*(µ, σ²) represents a normal distribution with a mean µ and variance σ².

Next, normalize simERP and simHEP amplitudes:

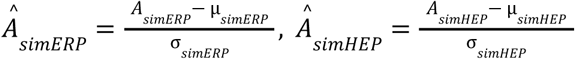

Calculate simHEPs which are related to simERP:

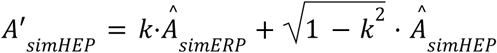

Where k is a scaling factor between 1 and 0 for positive and -1 and 0 for negative relationships. Rescale the resulting simHEP amplitudes:

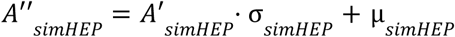

### 2.13. ΔECG with **Δ**EEG Correlation Analysis for Cardiac Field Artifact Influence

While there was no significant ECG difference between the investigated task conditions, we did observe significantly different ECG amplitudes between resting state and task recordings (Fig. 7C). Hence, it is possible that the observed HEP effects are explained by differences in cardiac field artifacts (CFA). CFA refers to the electric field generated by the heart which propagates through the body and is picked up by the EEG electrodes. To investigate the potential influence of the CFA on HEPs when comparing resting state and task recordings, we assessed whether the difference between the two conditions in ECG (Δ*ECG*) correlates to the difference in EEG (Δ*EEG*). Across all subjects, the correlation between Δ*ECG* and Δ*EEG* was assessed by Spearman correlation for each time point and electrode. Next, we identified clusters of significant correlations comprising a minimum of 2 neighboring channels. Surrogate statistics were calculated based on the correlation of randomly permuted Δ*ECG* time courses with the Δ*EEG*. This was the procedure repeated 1000 times with shuffled Δ*ECG*, each time retaining the summed *t*-value of the largest cluster. Lastly, we compared the summed *t*-values of each empirical cluster to the distribution of maximum summed *t*-values across all permutations. Clusters were considered significant if their summed *t*-value was larger than 95% of the summed *t*-values found in the surrogate analysis.

## 3. Results

### 3.1 HEP differences precede ERPs with high and low P300 amplitude

Given that HEP amplitudes were shown to be higher for interoceptive than exteroceptive attention (Al et al., 2021; Al, et al., 2020; García-Cordero et al., 2017; Petzschner et al., 2019; Villena-González et al., 2017) and P300 amplitudes index an attentional shift towards external stimuli (Kok, 2001; Polich, 2007) we hypothesized that HEPs should show a negative relationship with P300 amplitudes. In line with this prediction, we found significantly smaller HEPs preceding high P300 ERPs in the time between 156–600 ms in a posterior electrode cluster (Fig. 2A, cluster sum(*t*) = 2842, *p* ≤ 0.001, significant channels: CP6, TP9, TP10, P3, P4, P7, P8, Pz, O1, O2, PO9, PO10). This was reversed for frontal electrodes, with larger HEPs for larger P300 ERPs in a similar time window (time range: 144–600 ms, cluster sum(*t*) = -4562, *p* ≤ 0.001, significant channels: Fp1, Fp2, F3, F4, F7, F8, Fz, FC1, FC2, FC5, FC6, C3, C4, T7, Cz, CP5). The ECG was not significantly different between the high and low P300 conditions (temporal cluster-based permutation, *p* > 0.05), indicating that the effect is independent of the cardiac field artifacts.

**Figure 2.**
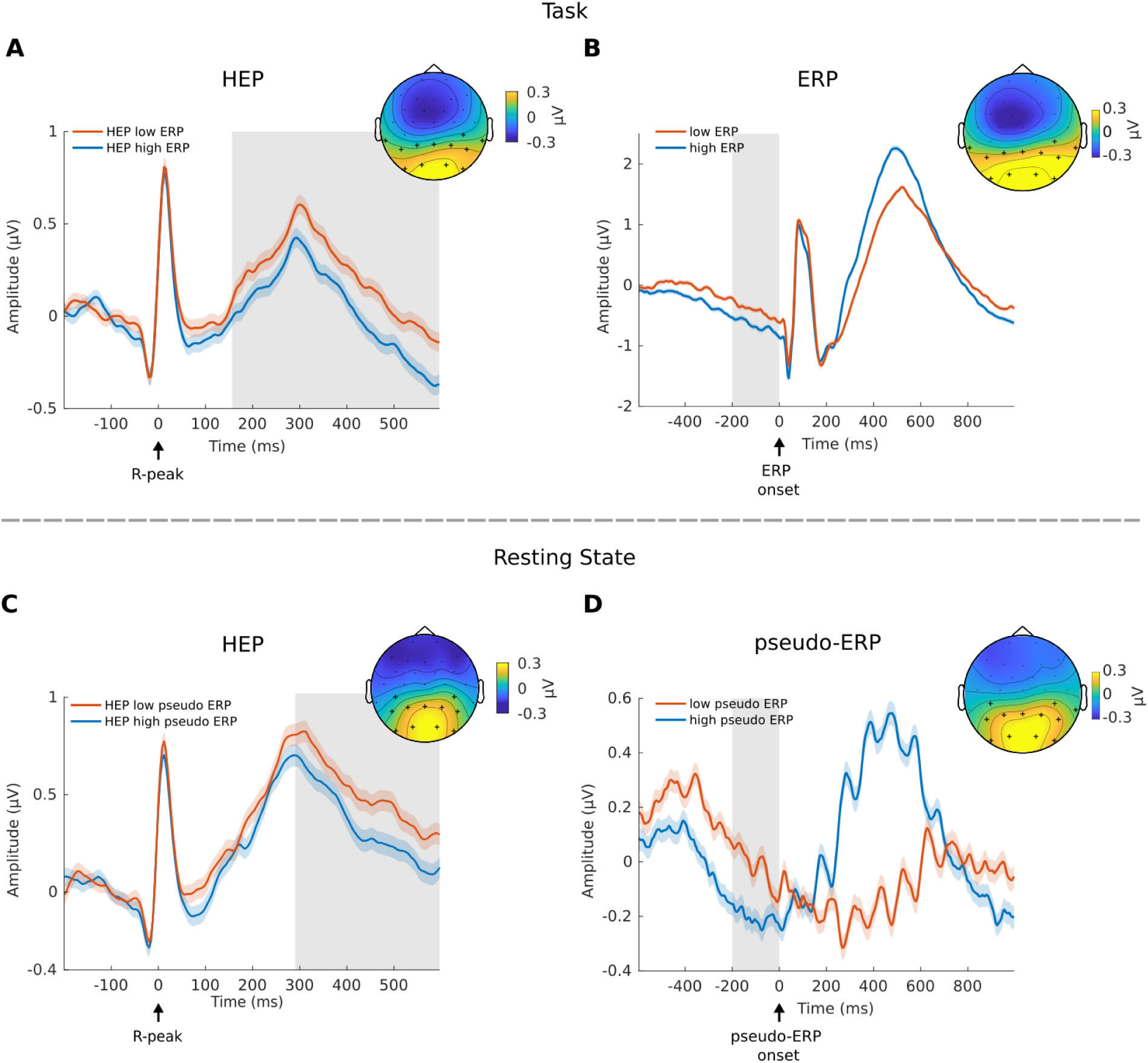
High amplitude P300 ERPs are preceded by low amplitude HEPs - also in the absence of a task. A) Pre-stimulus HEPs, categorized by the P300 amplitude of ERPs, reveal significantly higher HEP amplitudes preceding trials associated with low amplitude P300 ERPs, particularly over posterior electrodes. B) A comparison of pre-stimulus activity for low and high P300 amplitude ERPs, in the time window from -200 to 0 ms, also shows a significant difference. C) HEPs are significantly different if they are sorted by the amplitude of “pseudo-ERPs”. D) Resting state pseudo-ERPs show a significant difference in the time window of -200 – 0 ms. Shaded regions around the lines represent ± Standard Error of the Mean (SEM), grey boxes: *p*≤0.001, topoplots are averaged over time of significant differences and black crosses indicate significant channels.

However, when inspecting the sorted P300 ERPs, they show significant pre-stimulus baseline differences (Fig. 2B) (average between -200–0 ms, cluster-based permutation in space, cluster sum(*t*) = 121.5, *p* ≤ 0.001, significant channel: CP6, TP9, TP10, P3, P4, P7, P8, Pz, O1, O2, PO9, PO10). To further investigate whether this pre-stimulus difference can drive the observed HEP effect, we performed a surrogate heartbeat analysis. Within each participant, we created surrogates by permuting the original R-peak onset timings across trials per condition. Activity time-locked to the surrogate heartbeats was subsequently analyzed using cluster-based permutation testing while retaining the cluster *t*-values for the largest positive and negative clusters. This procedure was repeated 1000 times. The original *t*-values for the positive and negative cluster were not greater than the corresponding *t*-values obtained by the surrogate analysis (Monte-Carlo *p*-values positive cluster: *p* = 0.031, Fig. 3A, negative cluster: *p* = 0.249), indicating that the observed differences between HEPs in the high and low P300 condition are not time-locked to the R-peak.

**Figure 3.**
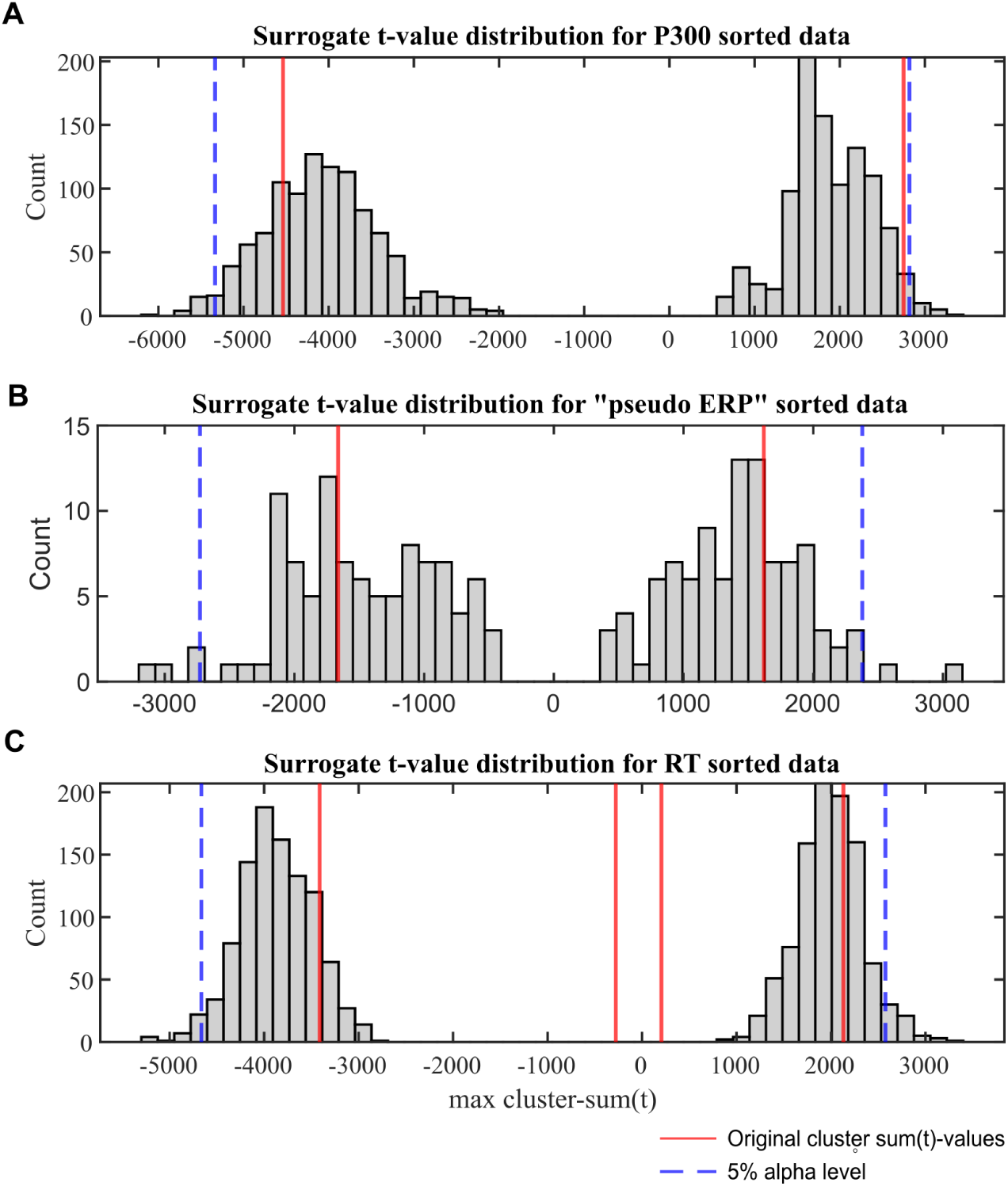
Distribution of cluster sum(*t*)-values obtained from surrogate heartbeat control analyses. Histograms show cluster sum(*t)*-values for surrogate heartbeat differences with shuffled R-peak onsets and original HEP effects for all significant clusters (red lines). The related trials are sorted in A) high vs. low P300 amplitude, B) high vs. low “pseudo ERP” amplitude, and C) fast vs. slow reaction time (RT). The dotted blue lines represent the 97.5% and 2.5% alpha cutoff.

### 3.2 During rest, HEP differences precede pseudo-ERPs with high and low amplitude in P300 time window

To determine whether the observed association between HEPs and P300 amplitudes is related to the data analysis approach, we performed a control analysis using resting state data. We copied the oddball task triggers from the task recording to resting state data of the same subjects and repeated the same analyses. Since during resting state, real HEPs are present, but task-evoked activity is absent, we can investigate if the HEP differences are task-related or a result of data-sorting. The resting state data was sorted into high & low amplitude conditions based on 250–600 ms post stimulus time window, creating high and low amplitude “pseudo-ERPs”, without any real task evoked activity (Fig. 2D) and HEPs that occurred in the pre-stimulus time of the “pseudo-ERPs” (Fig. 2C). The resting state control analysis included 1730 subjects and data processing was the same as used for real task data. Using spatio-temporal cluster-based permutation *t*-tests we observed smaller HEPs before trials with high “pseudo-ERP” amplitude between 292–600 ms in a posterior electrode cluster (Figure 2C, cluster sum(*t*) = 1618, *p* ≤ 0.001, significant channels: CP5, CP6, P3, P4, P7, P8, Pz, O1, O2, PO9, PO10). Negative clusters were present in the time 288–600 ms in a frontal electrode cluster (cluster sum(*t*) = -1663, *p* ≤ 0.001, significant channels: Fp1, Fp2, F3, F4, F7, F8, Fz, FC1, FC2, FC5, FC6, C3, T7, Cz, FT9, FT10; cluster 2: time range: 64 – 132 ms, sum(*t*) = -218, *p* = 0.014, significant channels: Fp1, Fp2, F3, F7, F8, FC5, FC6, C3, T7, FT9, FT10). In line with this, we again observed a significant difference in the pre-stimulus time -200-0 ms of the sorted resting state “pseudo ERPs” (Figure 2D, cluster sum(*t*) = -94, *p* ≤ 0.001, significant channels: CP5, CP6, TP10, P3, P4, P7, P8, Pz, O1, O2, PO9, PO10). These findings suggest that the HEP difference we observed when comparing high vs. low real (not “pseudo-ERPs”) P300 trials during the task is in fact independent of the task. Instead, it likely represents an artifact stemming from the sorting process of the trials, possibly due to regression towards the mean (Bland & Altman, 1994). This observation is further supported by a surrogate heartbeat control analysis, which showed that the sum(*t*)-values of HEP differences during rest was not bigger than the sum(*t*)-values of the surrogate data (Figure 3B, 100 permutations, positive cluster: *p* = 0.34, negative cluster *p* = 0.42)

### 3.3 HEP differences precede trials with fast and slow reaction times

The next aim of our study was to investigate whether HEP amplitudes relate to reaction times as a behavioral marker of task performance. Since reaction times relate to on-task attention (Posner & Boies, 1971; Yamashita et al., 2021), we hypothesized to see an inverse relationship with HEP amplitudes, with lower HEP amplitudes prior to faster reaction times. In addition, by using reaction times as a behavioral measure, we aimed to circumvent the artifacts induced by sorting post-stimulus data. We median-split trials in fast and slow RT categories and compared HEPs in the respective pre-stimulus time. Average reaction times were 432 ms (*SD* = 110 ms) for the fast condition and 529 ms (*SD* = 130 ms) for the slow condition. Based on spatio-temporal cluster-based permutation *t*-tests, we observed smaller HEP amplitudes for faster RT trials in two central electrode clusters (Figure 5A, negative cluster 1: 272–600 ms, cluster sum(*t*) = -3412, *p* ≤ 0.001, significant electrodes: F3, F4, Fz, FC1, FC2, FC6, C3, C4, Cz, CP5, CP6, P3, P4, Pz; negative cluster 2: 196–256 ms, cluster sum(*t*) = -277, *p* = 0.002, significant electrodes: FC1, FC2, FC6, C4, Cz, CP6, P4, Pz; positive cluster 1: 368–600 ms, cluster sum(*t*) = 204, *p* ≤ 0.001, significant electrodes: Fp1, Fp2, F7, F8, T7, T8, FT9, FT10, TP9, TP10, P7, O1, PO9, PO10; positive cluster 2: 320–356 ms, cluster sum(*t*) = 204, *p* = 0.019, significant electrodes: Fp1, Fp2, F7, F8, FC5, T7, FT9, FT10, TP9). The ECG was not significantly different between the fast vs. slow RT conditions (temporal cluster-based permutation, *p* > 0.05).

However, when closer inspecting the pre-stimulus time of the ERP trials split into a fast and slow RT category, it becomes apparent that the EEG activity before stimulus onset is already different between the two categories (Fig. 5B). Using spatial cluster-based permutation to compare the average amplitude in the -200–0 ms time we found a significant difference between the conditions in a central electrode cluster (cluster sum(*t*) = -163, *p* ≤ 0.001, significant electrodes: F3, F4, Fz, FC1, FC2, FC6, C3, C4, Cz, CP5, CP6, P3, P4, Pz) with more negative amplitude for faster RT trials. Since the potential is leading up to the target stimulus which required the subjects to press a button, and its specific spatial distribution, we consider it to be a contingent negative variation (CNV, Walter et al., 1964). To investigate if the pre-stimulus CNV difference between fast and slow RT trials is driving the HEP effects, we again performed a surrogate heartbeat analysis with permuted R-peak timings within each subject per condition. We created 1000 surrogate datasets with shuffled R-peak onsets. When comparing the original sum(*t*)-values to the surrogate sum(*t*)-values, we would conclude that the effects are not related to the R-peak with a *p* = 0.87 and *p* = 1 (negative clusters) as well as *p* = 0.33 and *p* = 1 (positive clusters, Fig. 4C).

**Figure 4.**
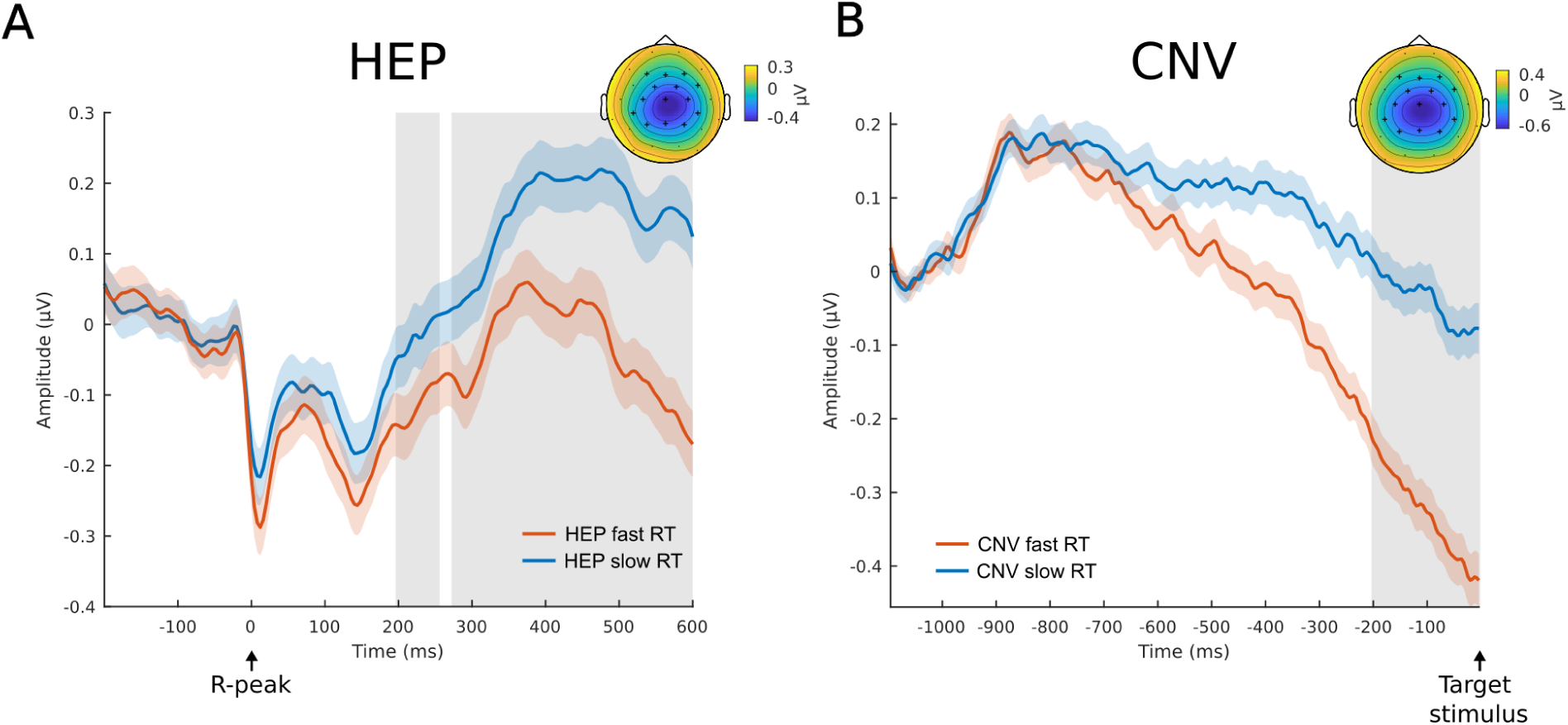
HEP and CNV Differences Before Fast and Slow Reaction Time Trials. A) HEP difference before fast and slow RT trials with significant differences in central electrodes. B) Difference in Contingency Negative Variation (CNV) before fast and slow reaction time (RT) trials. Pre-stimulus activity was compared between -200 – 0 ms and showed a significant difference. A baseline correction was applied based on the average amplitudes in the -1100 to -1000 ms time window. Grey area indicates *p* ≤ 0.001, Shades around the lines are ±SEM. Topoplots are averaged over time with the largest significant difference and significant channels are displayed as black cross.

### 3.4 No residual pre-stimulus HEP differences after pseudotrial correction

In the prior analyses, we demonstrated that the differences in pre-stimulus HEP observed when categorizing trials into either a high vs. low P300 condition or a fast vs. slow RT condition, are in part heartbeat unrelated. However, we wanted to investigate whether genuine HEP differences might still exist while being masked by the spurious effects we observed (see also simulation example 3.5). Hence, we’ve adopted a pseudotrial correction method (Huang et al., 2017; Wainio-Theberge et al., 2021), which subtracts ongoing fluctuations from the HEPs and can thereby correct for slow potential differences. We repeated our cluster-based permutation tests for the pseudotrial corrected data and did not observe any significant condition differences for the pre-stimulus HEPs between high vs. low P300

ERP amplitude, fast vs. slow reaction times, or pseudo-ERP amplitude during resting state (Fig. 6, cluster-based permutation *t*-tests, *p* > 0.05). Equivalence tests were conducted to investigate if the conditions can be considered statistically equivalent after pseudotrial correction. Per participant, we averaged the corrected HEP data across the electrode and time cluster which was significant before pseudotrial correcting the data. The effect size of the difference between conditions was then compared to the smallest meaningful effect of a Hedge’s *g* = 0.19 (see methods for explanation of this boundary). After correction, the pre-stimulus HEPs across all conditions can be considered equivalent, which implies that any initial significant differences between pre-stimulus HEPs were likely due to heartbeat independent confounding factors. This includes during the oddball task the high vs. low P300 (*t*_1739_ = -7.2, *p* < 0.01) and fast vs. slow RT (*t*_1739_ = 5.51, *p* < 0.01), as well as during resting state high vs. low pseudo-ERP (*t*_1730_ = -7, *p* < 0.01).

**Figure 5.**
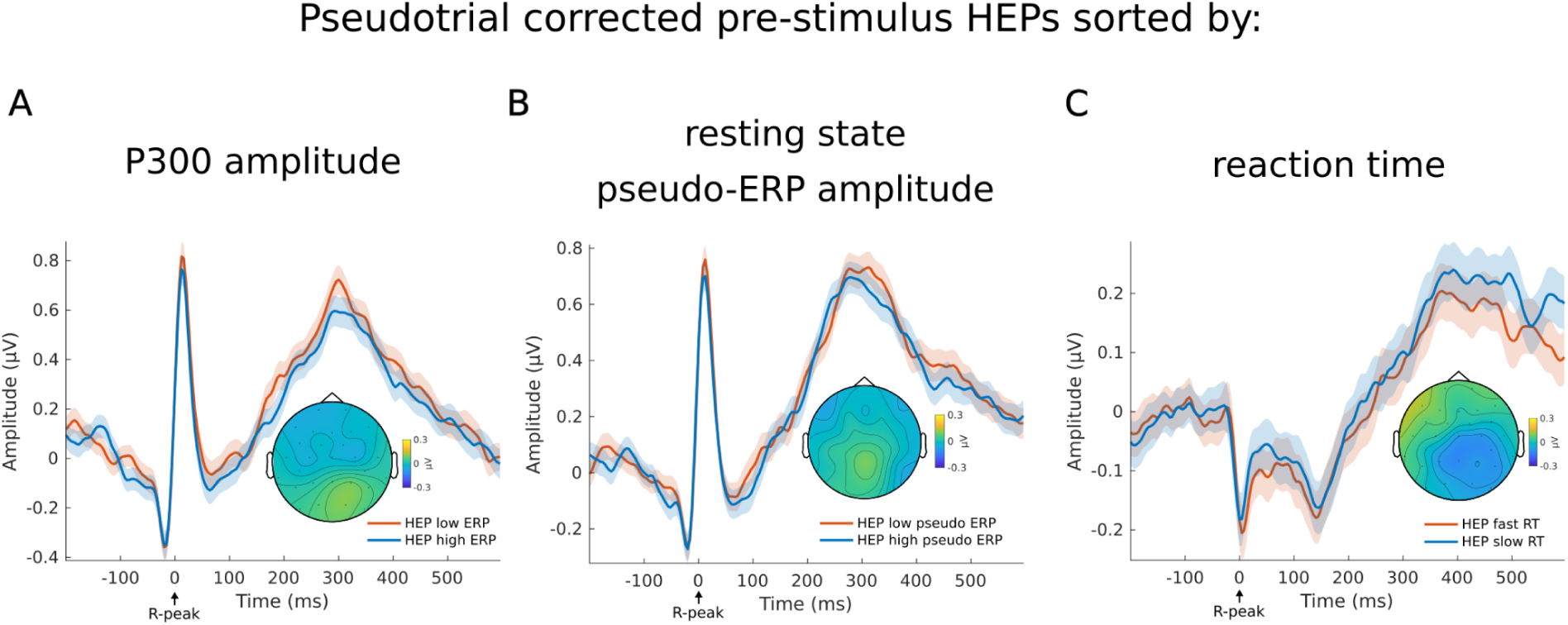
Pre-stimulus HEPs show no significant differences after pseudotrial correction. After pseudotrial correction, no significant differences can be found for pre-stimulus HEPs that were sorted based on A) the amplitude of the subsequent ERP P300 amplitude during the oddball task, B) the amplitude of pseudo-ERPs sorted in high vs. low condition in the P300 time window during resting state, or C) the reaction time speed during the oddball task. The average HEPs are based on the same electrode clusters used for the respective HEPs in Figure 1, 3 and 5. Shades around the lines are ± SEM.

**Figure 6.**
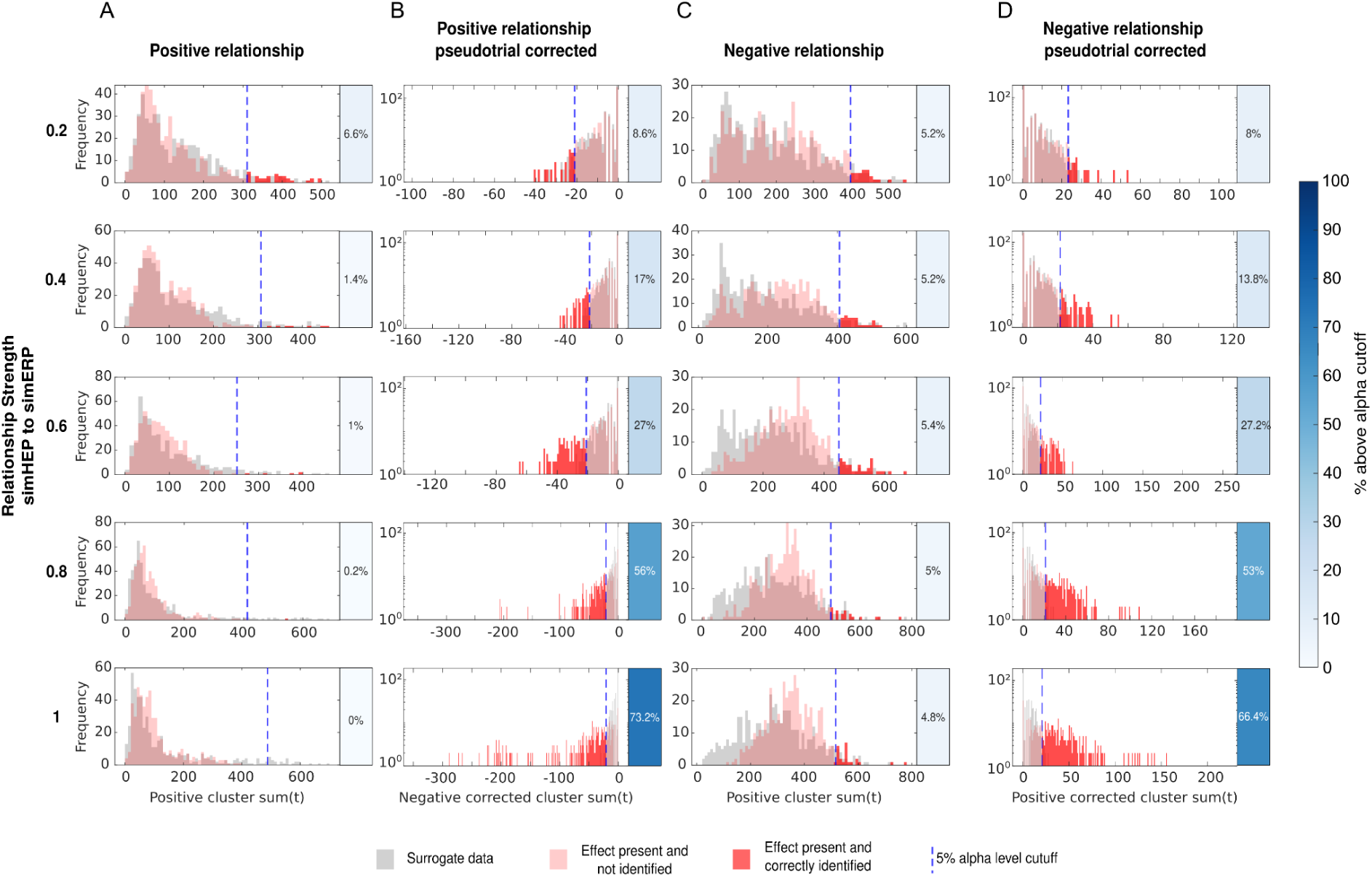
Pseudotrial correction and surrogate control analysis on simulated HEP to ERP relationships. Histograms visualizing cluster sum(*t*)-values obtained from 500 permutations of simulated data where an association between pre-stimulus simHEP and simERP was present (red) and corresponding surrogate data (gray). The dotted line indicates the 5% alpha level obtained from the surrogate data. sum(*t*)-values greater than this cutoff are considered significant in a surrogate heartbeat control analysis (dark red). Boxes right next to the histograms indicate which percentage of the permutations would be considered significant, and hence reflects the power to detect an effect. Rows relate to different linear relationship strengths between pre-stimulus HEP and post stimulus evoked response. Columns reflect positive or negative relationship between pre-stimulus HEP and post stimulus evoked response and their pseudotrial corrected counterparts. The y-axis of histograms after pseudotrial correction (B & D) is log-scaled for better visibility.

### 3.5 Removal of sorting-induced artifacts by pseudotrial correction increases sensitivity to detect true HEP effects in simulated data

Differences in HEPs between conditions can be spuriously introduced by overlap with ongoing brain activity (Park & Blanke, 2019). To account for this, surrogate heartbeat control analyses have been proposed. However, when the surrogate heartbeat procedure is used to control for the influence of non-heartbeat related activity, true HEP effects may be missed if they are mixed with confounds. Hence, we’ve adopted a pseudotrial correction method (Huang et al., 2017; Wainio-Theberge et al., 2021), which subtracts ongoing fluctuations from HEPs and has the potential to correct for artifacts. To demonstrate that pseudotrial correction can indeed uncover genuine HEP effects if they are mixed with other processes, we simulated EEG data where task evoked activity (simERP) relates to pre-stimulus HEP amplitudes (simHEP) in positive and negative direction (see methods for details). The simulated data was sorted based on the 250–600 ms post stimulus time window (corresponding to the P300 time range). Splitting data into high and low amplitude conditions can have long-lasting effects on the EEG, which can reach the pre-stimulus time (see Fig. 2 B/D) and induce spurious pre-stimulus HEP effects (see the preceding sections). To investigate if true simHEP effects can be identified in the presence of sorting induced artifacts, we performed a surrogate heartbeat control analysis on this data. The same analysis was repeated 500 times, while for each iteration the sum(*t*)-value of the largest cluster was retained for both: data where a relationship was present between simHEP and simERP (original data) but also their corresponding surrogate data. Based on the surrogate data, we defined a 5% alpha cutoff. Next, we calculated the percentage of permutations where the cluster sum(*t*) of the original data was larger than this cutoff. We found that significant HEP effects were detected in 0-6.6% of cases, demonstrating the effectiveness of surrogate heartbeat analysis in controlling for false positive findings in the presence of heartbeat independent confounds (type I error) (Fig. 6, A / C). However, even with a correlation of 1 between simHEP and simERP, in only 0% (positive relationship) or 4.8% (negative relationship) of the cases a genuine HEP effect would be found. Since real effects are present, but remain undetected (type II error) we performed a pseudotrial correction on the same data. We show that the correction can remove some of the spurious differences and uncover genuine simHEP effects, as the power to detect differences increased up to 73.2% (Fig. 6 B, positive relationship) and 66.4% (Fig. 6 D, negative relationship). It is important to note, that the effects were simulated with a conservatively small SNR of 3.3 dB for simERPs and -9.3 dB for simHEPs, which influences the ability to detect present effects.

### 3.6 HEP differences between oddball task and resting state

As shown in the previous sections, in our task, the pre-stimulus HEPs are overlapping with a slow stimulus preceding potential (CNV) and that these CNVs can effectively be removed from HEPs by pseudotrial correction. In simulations, we could show that slow artifacts can dominate HEP effects, which prevents other potentially present HEP effects from being found. Hence, our next aim was to investigate the relationship between resting-state and task HEPs before and after pseudotrial correction. We hypothesized that, due to the overlap with the CNV, HEPs would mirror its spatio-temporal distribution and appear lower in central electrodes during task than during rest prior pseudotrial correction. Furthermore, we hypothesized that pseudotrial correction would remove the spurious effect and uncover differences in HEPs between rest and task that are invisible before correction.

Before pseudotrial correction, we can only observe two clusters in positive as well as negative direction (Fig. 7A, positive cluster 1: 44-600 ms, cluster sum(*t*)=8046, *p* ≤ 0.001, positive cluster 2: -72–8 ms, cluster sum(*t*)=431, *p* ≤ 0.001; negative cluster 1: 24–600 ms, cluster sum(*t*): -8906, *p* ≤ 0.001; negative cluster 2: -72–20 ms, cluster sum(*t*): -375, *p* = 0.002). Surrogate heartbeat control analysis (100 iterations) indicates that these clusters are heartbeat unrelated, as all maximum sum(*t*) values obtained from shuffled data are larger than the original effects.

Importantly, the heartbeat independent effect can be removed by pseudotrial correction, revealing several clusters distributed in time and space (the first two clusters with the largest sum(t) values are illustrated in Fig. 7B, for a summary of all clusters see supplementary Table 2). Surrogate heartbeat control analysis indicates that all clusters in positive as well as negative direction have sum(*t*) values larger than 97.5% or smaller than 2.5% of the cluster sum(*t*) values obtained from shuffled data. Therefore, we can conclude that the effects we observe after pseudotrial correction reflect differences in activity which are time-locked to the heartbeat.

To assess the potential influence of cardiac field artifacts on these differences, we compared the ECG between resting state and task by a temporal cluster-based permutation *t*-test. Several significant differences could be observed (Fig. 7C, *p* ≤ 0.001), indicating possible CFA induced HEP differences when comparing resting state and task. Furthermore, differences in heart rate were present, with 61.3 (*SD* = 8.1) beats per minute (bpm) during rest and 63.7 (*SD* = 8.2) bpm during the task (paired *t*-test, *p* = 1.0828e-39). To further investigate if the CFA can explain the HEP differences, we tested whether the difference between resting state and task ECG (Δ*ECG*) correlates to the difference between resting state and task EEG (Δ*EEG*). Yet no significant clusters were found in the HEP time window of interest (Fig. 8).

**Figure 7.**
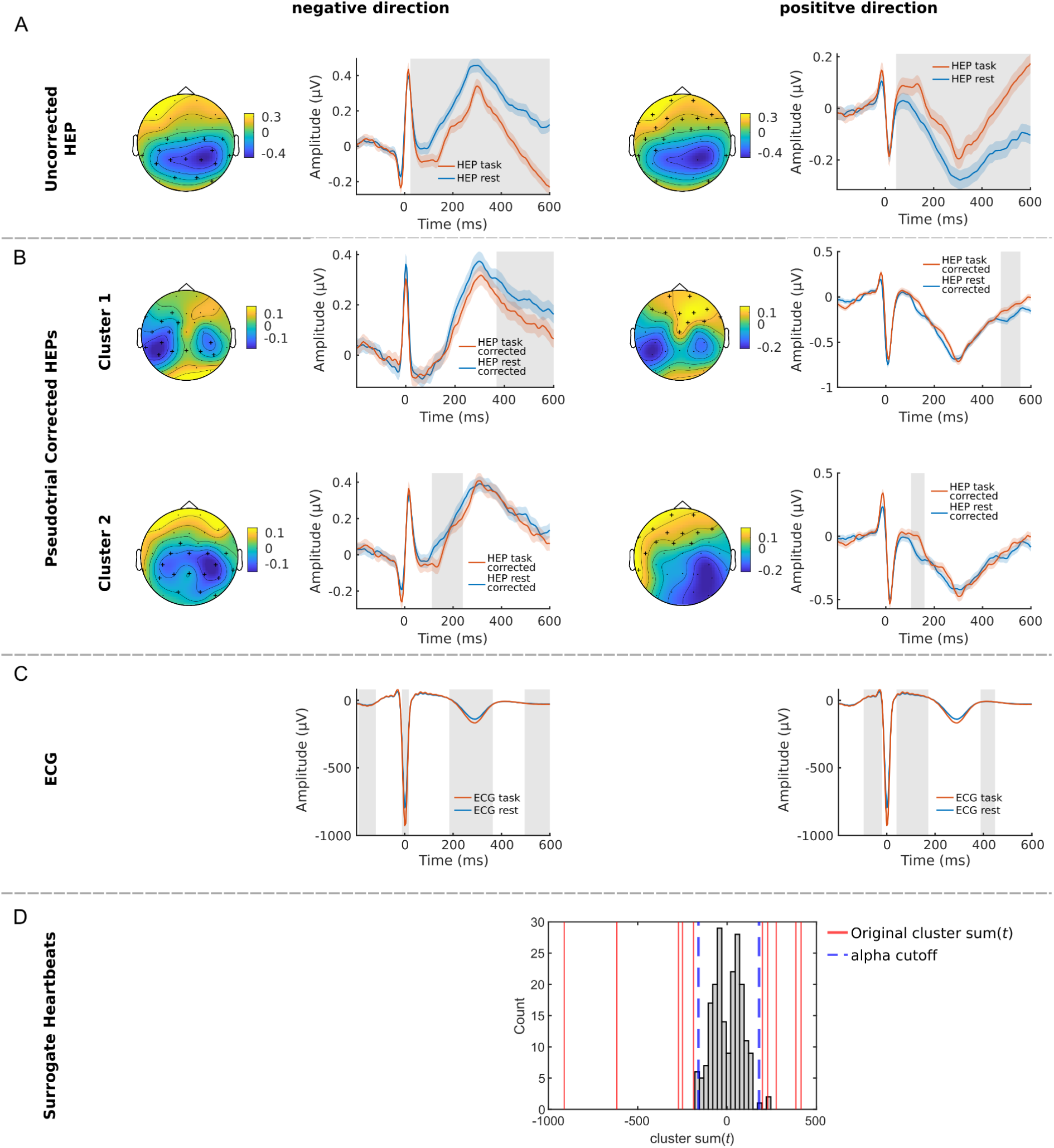
HEP differences between Oddball task and resting state. Differences in HEPs between Oddball task and resting state in positive and negative direction A) before and B) after pseudotrial correction. After correction, several clusters were found and the largest first two are illustrated. C) ECG differences between task and rest are illustrated. Shades around the lines are ±SEM, grey boxes: *p* ≤ 0.05, Topoplots are averaged over time of significant differences and black crosses highlight significant channels. D) Histograms visualizing the surrogate heartbeat control analysis. For positive as well as negative differences, all clusters of the original data have larger summed *t*-values (red lines) than the 95% or smaller than the 2.5% alpha cutoff (blue dashed lines) obtained from the largest surrogate heartbeat summed *t*-values (gray bars).

**Figure 8.**
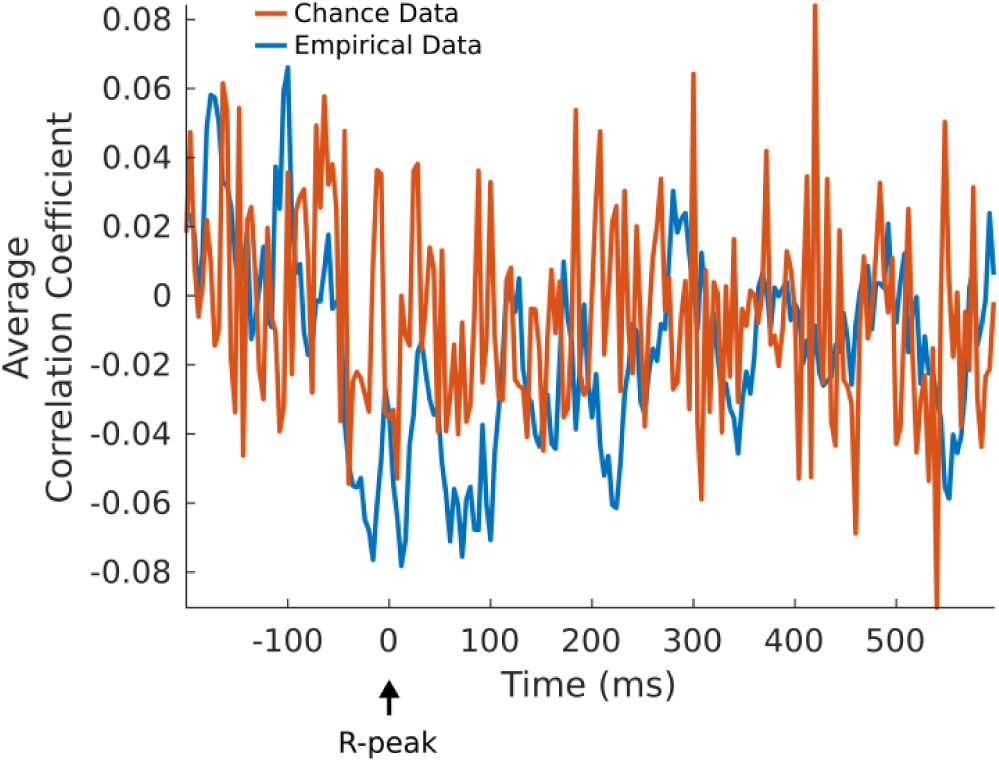
No Significant Correlation between Δ*ECG* and Δ*EEG*in time range of HEP analysis. Correlation between resting-state and task-related differences in ECG (Δ*ECG*) and EEG (Δ*EEG*) across all channels and time points and corresponding chance data obtained by correlation of Δ*EEG* with randomized Δ*ECG* data. No clusters of significant differences were found (*p* > 0.05). Time courses for illustration are obtained from the Cz electrode and grand averaged across subjects.

**Figure 9.**
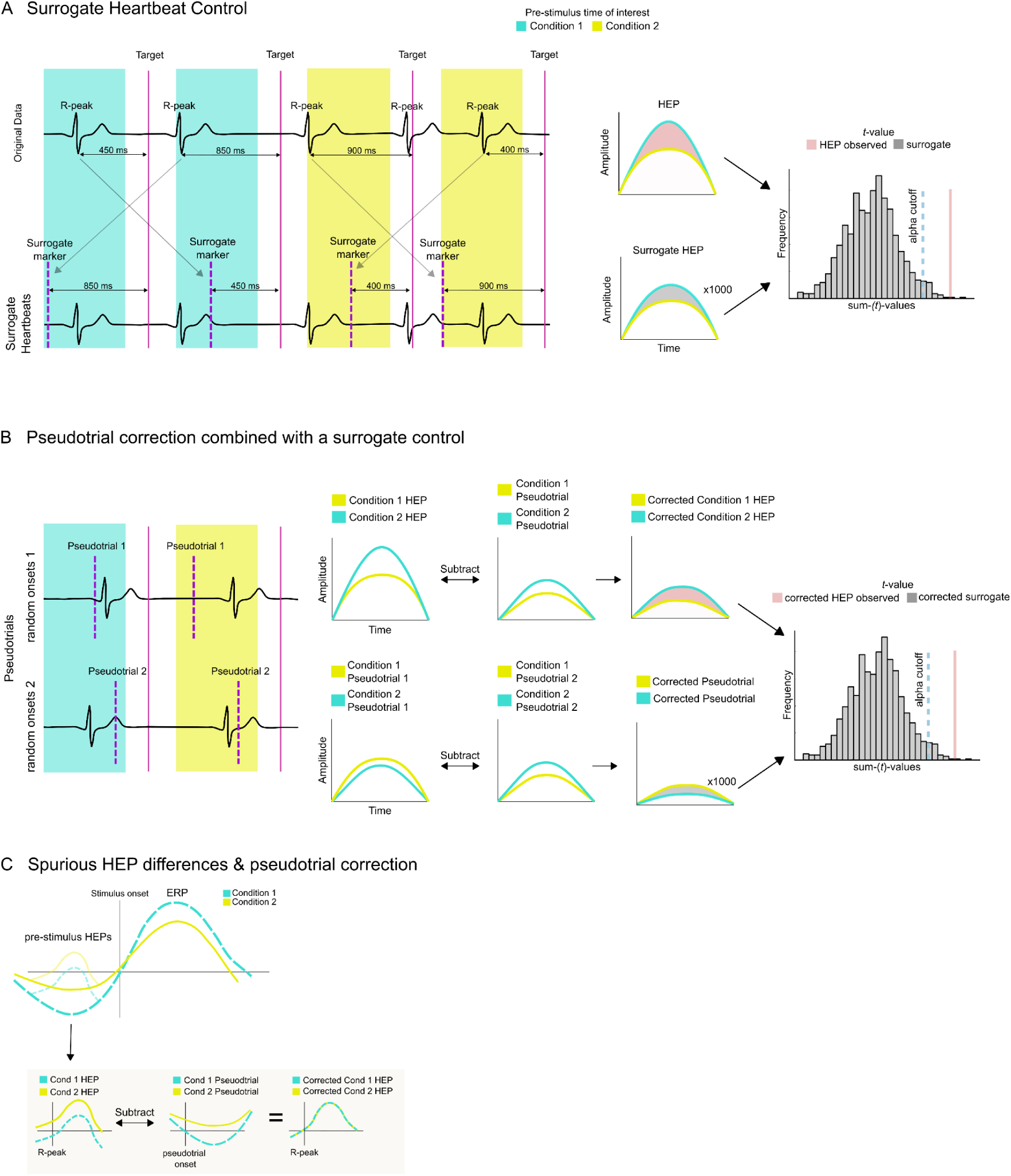
Schematic overview of the main methods used in this study. A) Illustration of the surrogate heartbeat procedure adapted in this study. HEPs are calculated based on heartbeats occurring the pre-stimulus time of two different conditions. To control for heartbeat unrelated effects, R-peak timings relative to the target stimuli are shuffled within each condition, generating surrogate HEPs. This procedure is repeated a number of times, each time calculating the condition difference between surrogate HEPs. The resulting null-distribution of cluster sum(*t*)-values is used to assess the significance of the original HEP effect. B) Pseudotrial are generated by adding random triggers to the pre-stimulus time of interest. HEPs are corrected by subtracting pseudotrials per condition, potentially revealing some residual effect. To control effects introduced by the pseudotrial correction, different permutations of random pseudotrials are subtracted from each other per condition. This procedure is repeated a number of times, each time calculating the condition difference between the corrected pseudotrials. The resulting null-distribution of cluster sum(*t*)-values is used to assess the significance of the original HEP effect. C) Schematic representation of spurious HEP differences based on heartbeat unrelated condition differences. Subtraction of pseudotrials removes unrelated activity.

## Discussion

Heartbeat evoked potentials, which reflect the cortical processing of afferent signals generated by the heartbeat, have been shown to relate to different cognitive, emotional, or pathophysiological conditions (Engelen et al., 2023; Park & Blanke, 2019). However, when investigating HEPs in different conditions, it is important to control for potential overlaps with ongoing neural activity unrelated to the heartbeat (Azzalini, et al., 2019). In the present study, we advocate for the use of appropriate control analyses, such as pseudotrial correction (Wainio-Theberge et al., 2021) and surrogate heartbeat analyses (Azzalini, et al., 2019; Park & Blanke, 2019) to account for heartbeat unrelated confounds. In addition, in simulated and real data we show that pseudotrial correction has the potential to remove some of the heartbeat unrelated confounds and uncover genuine HEP effects.

### Pre-stimulus HEPs effects can be spuriously induced by underlying differences in neural activity

It was previously proposed that HEP amplitudes may serve as a marker for attentional processes - with higher amplitudes being indicative of interoceptive attention, and lower amplitudes for exteroceptive attention (Al, et al., 2020; Petzschner et al., 2019). Given that P300 ERP is a signature of task relevant attentional processes (Kok, 2001; Polich, 2007), we expected P300 amplitudes to be inversely related to pre-stimulus HEP amplitudes. To test for this, we sorted single trial P300 ERPs into high and low amplitude subgroups and investigated whether there is a significant difference between the corresponding pre-stimulus HEPs. Initially, we indeed find the expected significant differences. However, when performing the same analysis on resting-state data, generating high and low “pseudo ERP” conditions, we could replicate a similar observation. This finding was surprising, since during rest no task was performed, and we hence would not expect a meaningful effect of high vs low amplitude states on HEPs.

To investigate the origin of the observed HEP differences further, we first tested whether the effects we find are actually related to the heartbeat. For this, we created surrogate heartbeats by shuffling the original timings of the HEP onsets. Using the resulting surrogate HEPs, we then repeated the same statistical analysis as described above many times, generating a null-distribution of effects. Comparing our original results to the surrogate null-distribution, the original HEPs differences could not be considered significant. This indicates that the initial HEP effects were likely unrelated to the heartbeat.

Thus, the question arises why we initially observed differences in pre-stimulus HEPs? One potential explanation might lie in the inherent temporal dependency of the EEG signal, where the up- and down-states of the signal do not fluctuate randomly, but are organized in a sequence over time. Due to the cyclic nature of the EEG signal, when investigating the association between high amplitudes in the post-stimulus time (up state), it is likely that they are preceded by low amplitudes in the pre-stimulus time (down state) and vice versa. This can creates an inverse correlation between the two time-points. Due to the relatively long time range spanning pre- and post-stimulus time in our experiment, especially slower fluctuations will shape the relationship between the two. These fluctuations are characterized by oscillations in the low frequency < 4 Hz range (e.g. Delta rhythm and below), but also faster oscillations can influence their dynamics on longer time-scales via a baseline shift mechanism (Studenova et al., 2023) or through phase amplitude coupling (Tort et al., 2010). In addition, regression to the mean, where extreme values at one time-point are followed by less extreme values at another time-point (Bland & Altman, 1994; Tu & Gilthorpe, 2007), likely plays a role in the inverse relationship between HEP and ERP amplitude.

However, even though the HEP differences we observe are likely unrelated to the heartbeat, the possibility still exists that genuine HEP effects are present, but not found because the spurious activity is too strong. We accounted for this by performing a pseudotrial subtraction which effectively subtracts heartbeat-unrelated activity. After applying this correction, the initial HEP differences could not be considered significant anymore, indicating that they were likely due to confounding neural activity. Our results are in contrast to previously reported inverse relationships between pre-stimulus HEPs and P300 amplitudes (Al et al., 2021; Al, et al., 2020; Marshall et al., 2019). While differences in the used tasks (somatosensory detection or reward incentive paradigm) and analysis strategies are apparent, further research is needed to elucidate the scenarios in which a relationship between pre-stimulus HEPs and the P300 ERP is further validated.

It is important to note that not only the investigation of different amplitude conditions can be problematic. In our second example, we show that by using the behavioral variable reaction time, we can create a similar situation. We hypothesize that pre-stimulus HEPs are related to reaction times, since reaction time speed is related to on-task attention (Posner & Boies, 1971), and HEPs are lower if the focus of attention is on task-related processes (Al et al., 2020; Petzschner et al., 2019; Zaccaro, 2022). While we initially observed lower HEPs with faster reaction times, our pseudotrial correction and surrogate heartbeat analysis uncovered their spuriousness. One reason for the spurious effect is a a slow preparatory potential in the pre-stimulus time of target stimuli which is also related to reaction times. It likely reflects a contingent negative variation (CNV), which is related to anticipatory attention and response preparation (Hillyard, 1969; Walter et al., 1964). Plotting the initially significant HEP differences, the topoplot looks strikingly similar to the CNV. Hence, after sorting trials based on reaction times, the CNV is confounding the HEPs, with more CNV influence in fast RT trials, and less CNV in slow RT trials. Notably, a recent study that used a similar correction method—subtracting pseudotrials from HEPs in the presence of CNV—and also found no association between HEP amplitudes and anticipatory attention (Aprile et al., 2024 - preprint). Consistent with this, (Marshall et al., 2019) observed no direct association between pre-stimulus HEP and CNV amplitude or reaction times. Our results are therefore in line with literature and provide no evidence for an interaction between reaction times and pre-stimulus HEP amplitudes.

### The potential of pseudotrial correction to uncover genuine HEP effects

Overall, after carefully controlling for confounding factors, there appear to be no significant HEP effects in our task based analyses. However, we wanted to illustrate the potential of pseudotrail correction to uncover small effects which are present, but are missed due to the overlap with stronger heartbeat-unrelated activity. For this, we simulated data where pre-stimulus HEPs (simHEP) are related to post-stimulus ERPs (simERP) in positive and negative direction. As in the previous analysis, we sorted the simERPs into different amplitude conditions, and investigated whether pre-stimulus simHEPs would differ significantly among conditions. Before pseudotrial correction, most HEP differences would be considered heartbeat independent based on a surrogate procedure. Importantly, after applying pseudotrial correction, the probability to correctly identify HEP effects increases with the strength of the relationship between simulated simHEP and simERP. This demonstrates that pseudotrial correction effectively mitigates the influence of artifacts and allows for a more accurate identification of true HEP effects.

### Pseudotrial correction can uncover previously undetected HEP effects in real data

To further show the potential of pseudotrial correction to remove spurious activity in real (not simulated) data, we compared HEPs obtained from resting state data with HEPs obtained from the pre-stimulus time of the oddball task. Because HEP amplitudes are higher when the attention is focussed on internal stimuli (Al, et al., 2020; García-Cordero et al., 2017; Marshall et al., 2017; Petzschner et al., 2019), their amplitude could be higher during rest compared to task (Al et al., 2021). In the task-based analyses, we could show that the pre-stimulus HEPs are overlapping with CNV, while during resting-state, this should not be the case. Because the CNV is a “downward drift” and HEP “rides on top of that”, we expect to find HEP amplitude which are spuriously lower during task as compared to rest. This was indeed the case, which was also supported by a surrogate heartbeat control analysis, indicating that much of the observed task vs. rest HEP difference was unrelated to the heartbeat. Notably, after pseudotrial correcting the HEPs, the initially present strong differences disappeared, revealing multiple smaller clusters with varying temporal and spatial distributions. These clusters stayed significant, even after application of surrogate control analysis, suggesting that they are likely heartbeat-related. It hence seems that pseudotrial correction successfully removed the CNV related confound and uncovered heartbeat locked HEP effects.

While the surrogate heartbeat and pseudotrial correction control for heartbeat-unrelated confounds, artifacts time-locked to the heartbeat can still play a role. To investigate if cardiac field artifacts (CFA; Dirlich et al., 1997), which represent the influence of the heart’s electrical activity on EEG signals, could affect the HEP results, we compared the ECG between resting-state and task conditions. Indeed, we observe several clusters of significant differences in the ECG between resting state and task. To assess whether the HEP differences between rest and task were solely driven by the CFA, we performed an additional control analysis. We first computed the differences between rest-task ECG (Δ*ECG*) and rest-task EEG (Δ*EEG*). We then performed a cluster-based correlation analysis across all time-points and channels between Δ*ECG* and Δ*EEG*. Since no cluster of significant correlations within the time-range of our HEP analysis was found (-200 to 600 ms around the R-peak), the analysis suggests that the observed HEP effects are not attributable to CFA only.

Overall our analysis therefore highlights the potential of pseudotrial correction to uncover otherwise hidden effects. Importantly, surrogate heartbeat control analysis can be applied to pseudotrial corrected data to account for residual noise or condition differences introduced by the pseudotrial subtraction. Therefore, by combining both methods, an effective control for heartbeat independent type I and II errors is achieved.

### Implications for Future Research on HEPs

The theoretical implications of slow potential differences in the ongoing neural activity on HEP amplitudes have been highlighted in prior work (Azzalini, et al., 2019; Park & Blanke, 2019). Importantly, many studies already incorporate surrogate heartbeat analyses to control for potential false positive findings (e.g. a non-exhaustive list Azzalini et al., 2021; Babo-Rebelo et al., 2016; Marshall et al., 2017, 2022; Park et al., 2014; Perogamvros et al., 2019; Simor et al., 2021). Our study demonstrates that, while this method effectively controls for false positives related to non-heartbeat confounds, genuine HEP effects can be missed if confounds are present. Therefore, combining pseudotrial correction and surrogate heartbeat analyses offers a solution to the problem of overlapping activity. For instance, Tanaka et al. (2023) report that their observed HEP effects may be a result of the overlap between task-evoked activity and their studied HEPs. Here, application of the correction methods could help to delineate the true task effect on HEPs.

Another topic of discussion is baseline correction in the field of HEP research. As previously suggested, due to the cyclic nature of the heartbeat, baseline correction can be problematic as the baseline window is not necessarily free of activity originating from the previous heartbeat (Azzalini, et al., 2019; Banellis & Cruse, 2020; Kern et al., 2013; Park et al., 2014; Petzschner et al., 2019). To address this concern, we performed all our main analyses with and without applying baseline correction. In our study, the observation of no pre-stimulus HEP effects in relation to P300 or reaction time were consistent regardless of baseline correction (see Fig S1-4). However, we note that the results were also not exactly the same. When no baseline correction is applied, resting state and task HEPs were shifted, which is corrected to some degree by pseudotrial correction (Fig S6). Also, after pseudotrial correction in the rest vs task comparison, a HEP difference appeared in the pre-stimulus time window in a fronto-central electrode cluster (Fig. S2, Cluster 1 in negative direction). When baseline correction is performed, this effect disappears, while potentially shifting post-stimulus HEP in a positive direction (Fig. 7). This example highlights the need for careful evaluation of the baseline window if baseline correction is performed (Luck, 2005).

### Limitations

Firstly, the population we used consisted of an elderly sample aged between 40 and 80 years old. Since previous research has shown that resting state HEPs are affected by aging (Kamp et al., 2021), it could be that our null-findings are specific to this age group. However, others have specifically examined potential interactions between young and old age groups with HEPs during different cognitive tasks and observed no effects of age (Aprile et al., 2024 - preprint).

Our comparison of task vs. rest HEPs illustrates the potential of pseudotrial correction to uncover effects that may remain undetected in uncorrected data. However, in our experiment, also differences in ECG and heart rate were present which can potentially lead to spurious HEP effects. Although our correlation control analysis did not indicate a strong influence of the CFA on HEP results, some limitations should be acknowledged. We only have a single ECG channel available, and previous studies have shown that condition-induced ECG changes can vary depending on ECG electrode placement (Gray et al., 2007). It is still possible that the changes in HEP we observe are a reflection of the CFA, but that EEG and ECG electrodes capture different orientations of this artifact. Understanding how the CFA translates to EEG signals is crucial and should be addressed in future HEPs studies, ideally by using multiple ECG electrodes to provide a broader coverage of the cardiac artifacts. Therefore, it is essential that our results are replicated in carefully controlled experiments where cardiac and other physiological parameters are kept constant across conditions.

The surrogate heartbeat procedure generates a distribution of surrogate test statistics, which is compared to data obtained from real heartbeats. Since the resulting distribution is essential to make reliable comparisons, it has to be performed many times to serve as an effective control. However, the optimal number of iterations is not yet empirically established, and researchers choose a number between 100 (Babo-Rebelo, Richter, et al., 2016; Candia-Rivera et al., 2023; Engelen et al., 2022; Perogamvros et al., 2019), 200 (Park et al., 2016), 400 (Marshall et al., 2017), 500 (Azzalini, et al., 2019), and 1000 (Babo-Rebelo et al., 2019; Babo-Rebelo, Wolpert, et al., 2016) iterations. In our study, we selected different numbers of permutations for different analyses, primarily because the maximum number of feasible iterations is limited by available computational power, since performing spatio-temporal cluster-based permutation tests many times is computationally intensive. However, if effects are close to the significance threshold, it is advisable to increase the number of permutations to get a more accurate *p*-value estimate (Good, 1994).

Lastly, regression based ERP analyses are becoming more popular and offer a solution to the problem of overlapping ERPs (Burns et al., 2013; Ehinger & Dimigen, 2019; Smith & Kutas, 2015). In theory, pseudotrials can be added to the regression models as confounders to allow for the correction of heartbeat-unrelated activity. This form of pseudotrial correction needs to be established in future studies.

### Conclusion

Our study highlights the issue that neural activities which are not triggered by the heartbeat can interfere with the HEP if they occur at the same time. While many studies investigate HEPs during tasks, it is essential to distinguish HEP from other task-related activations, such as stimulus evoked responses or response preparation. In such experimental designs, even if careful statistical analysis are applied that incorporate different sources of confounders (e.g. cardiac and other physiological parameters), HEPs can still be confounded by heartbeat unrelated factors. We believe that careful experimental design, which avoids the overlap between heartbeats and task-related activities, in addition to adequate control analyses should be taken into account when investigating HEPs.

In this study we highlight the use of two control analyses which, when combined, represent a methodological advancement in the analysis of HEPs. By effectively isolating genuine HEP effects from confounding activity, these methods allow for more accurate interpretations of the interplay between neural processing of cardiac information and external stimuli.

## Data & Code availability

Anonymized data can be requested by application at the LIFE-Study Department (https://ldp.life.uni-leipzig.de/). The code used for the analyses will be made available at https://github.com/PaulSteinfath upon publication of the manuscript.

## Acknowledgements

This study has been supported by the LIFE-Leipzig Research Center for Civilization Diseases, Leipzig University. LIFE is funded by means of the European Union, by the European Regional Development Fund (ERDF), by the European Social Fund (ESF) and by means of the Free State of Saxony within the framework of the excellence initiative. We would like to thank all study participants and the LIFE team who made this study possible.

## Competing interests statement

The Authors declare no competing interests in relation to the work.

## Notes

### Competing Interest Statement

The authors have declared no competing interest.

