## Supplementary Materials for "Validating genuine changes in Heartbeat Evoked Potentials using Pseudotrials and Surrogate Procedures"

Due to the cyclic nature of the heartbeat and therefore potential overlap of the baseline correction window with the previous heartbeat, we chose to repeat all our main analyses without performing a baseline correction.

### **S1 HEP differences precede ERPs with high and low P300 and “pseudo-ERP” amplitude**

In line with the results obtained from baseline corrected data, we observe significantly different HEPs before trials sorted into high and low P300 amplitude conditions during an auditory oddbal task. Several significant clusters can be observed: positive cluster 1: sum(*t*): 2930, p ≤ 0.001, time: 156–600 ms, channels: T8, FT10, CP6, TP9, TP10, P3, P4, P7, P8, Pz, O1, O2, PO9, PO10; negative cluster 1: sum(*t*): -4536, p ≤ 0.001, time: 140–600 ms, channels: Fp1, Fp2, F3, F4, F7, F8, Fz, FC1, FC2, FC5, FC6, C3, C4, T7, Cz, FT9, CP5, negative cluster 2: sum(*t*): -345, p ≤ 0.001, time: 28–132 ms, channels: Fp1, F3, F7, Fz, FC1, FC2, FC5, C3, Cz, CP5. The largest positive cluster is visualized in Supplementary Figure 1A. Both clusters reflect a heartbeat independent process after the application of surrogate heartbeat analysis (SFig. 3A, positive-cluster: *p* = 0.1, negative-cluster: *p* = 0.9).
Furthermore, also if the same analysis is repeated after copying task triggers to resting state data a similar picture emerges with several significantly different clusters: positive cluster 1: sum(*t*): 2691, p ≤ 0.001, 176–600 ms, channels: CP5, CP6, P3, P4, P7, P8, Pz, O1, O2, PO9, PO10; positive cluster 2: sum(*t*): 384, time: 56–152 ms, ,p ≤ 0.001; channels: CP6, P3, P4, P7, P8, Pz, O1, O2; negative cluster 1: sum(*t*): -3005, p ≤ 0.001, 256–600 ms, channels: Fp1, Fp2, F3, F4, F7, F8, Fz, FC1, FC2, FC5, FC6, C3, C4, T7, Cz, FT9, FT10, TP9; negative cluster 2: sum(*t*): -343, p ≤ 0.001, 72–156 ms, channels: Fp1, F3, F7, FC1, FC2, FC5, C3, T7, Cz, FT9. The largest positive cluster is visualized in Supplementary Figure 1B. All clusters reflect heartbeat independent processes after the application of surrogate heartbeat analysis (SFig. 3B, positive-cluster: *p* = 0.59, negative-cluster: *p* = 0.7).

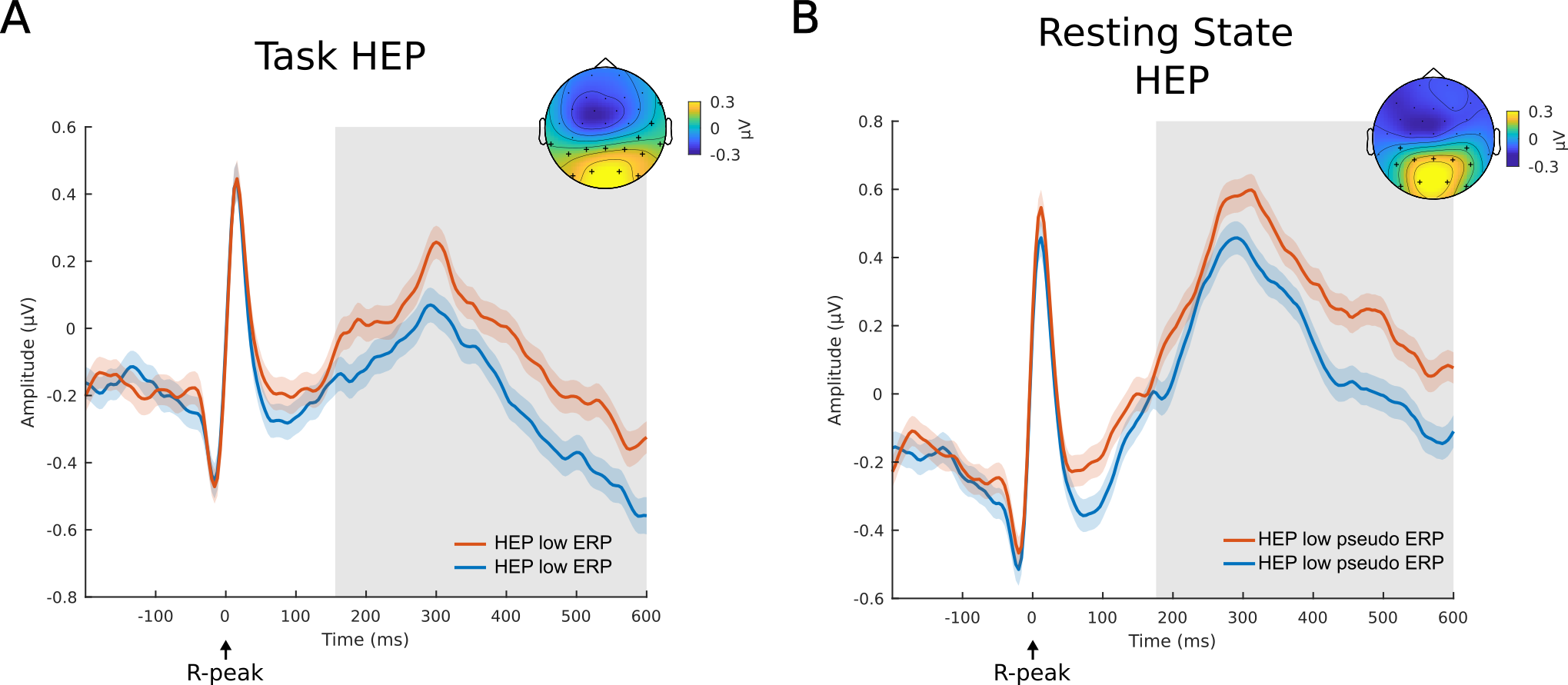

#### **Supplementary Figure 1. HEP differences precede trials sorted in high and low amplitude conditions during task and resting state**

A) HEPs preceding target trials sorted into high and low P300 ERPs are significantly different. B) During resting-state, HEPs sorted by the amplitude of “pseudo-ERPs” are significantly different. For resting state and task HEPs the largest positive clusters are shown. Shaded regions around the lines represent ±SEM, gray boxes: p≤0.05, topoplots reflect the average over the largest cluster of significance with black crosses indicating significant channels.

### **S2 HEP differences precede ERPs with fast and slow reaction times**

In line with the results obtained from baseline corrected data, we observe significantly different HEPs before trials were sorted into fast and slow reaction time conditions: negative cluster 1: sum(*t*) = -1494, p ≤ 0.001, time: 388 – 600 ms, electrodes: Fz, FC1, FC2, C3, C4, Cz, CP5, CP6, P3, P4, Pz; negative cluster 2: sum(*t*) = -220, *p* = 0.012, time: -124 – -80, electrodes: Fp1, Fp2, F7, F8, FC5, T7, FT9, FT10, TP9) and positive cluster 1: sum(*t*) = 818, p ≤ 0.001, time: 464 – 600 ms, electrodes: Fp1, Fp2, F7, F8, FC5, T7, FT9, FT10, TP9; cluster 2: sum(*t*) = 610, p ≤ 0.001, time: -196 – -72 ms, electrodes: F3, Fz, FC1, FC2, FC5, C3, C4, Cz, CP5, CP6, P3, P4, P8, Pz, O2. The largest negative cluster is visualized in supplementary figure 2A. All clusters reflect heartbeat independent processes according to a surrogate heartbeat analysis (SFig. 3C, negative cluster 1: *p* = 0.92, negative cluster 2: *p* = 1; positive-cluster 1: *p* = 0.85, positive-cluster 2: *p* = 0.98).
Furthermore, the pre-stimulus time (-200-0 ms) of target trials was significantly different with a similar spatio-temporal distribution to the HEPs (SFig. 2B, cluster sum(*t*) = -185, p ≤ 0.001, electrodes: F4 , Fz , FC1 , FC2 , FC6 , C3 , C4 , Cz , CP5 , CP6, P3 , P4 , Pz).

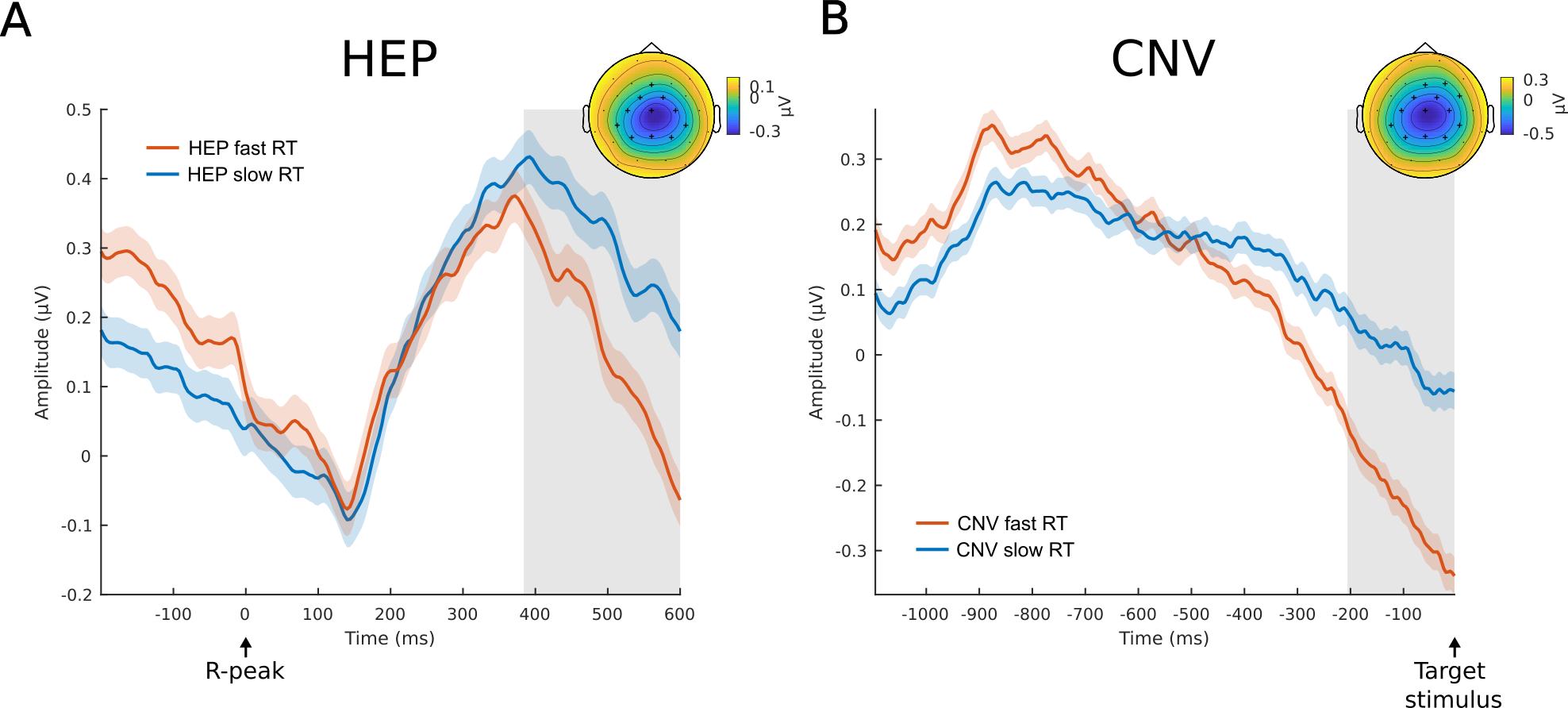
**Supplementary Figure 2 pre-stimulus HEPs sorted by RT during task**

A) HEP difference before fast and slow reaction time (RT) trials with a late significant negative cluster between 388–600 ms in central electrodes. B) Difference in Contingent Negative Variation (CNV) before fast and slow RT trials. No baseline correction was applied. Shades around the lines are ±SEM. gray boxes: p≤0.05, topoplots reflect the average over largest cluster of significant difference with black crosses indicating significant channels.

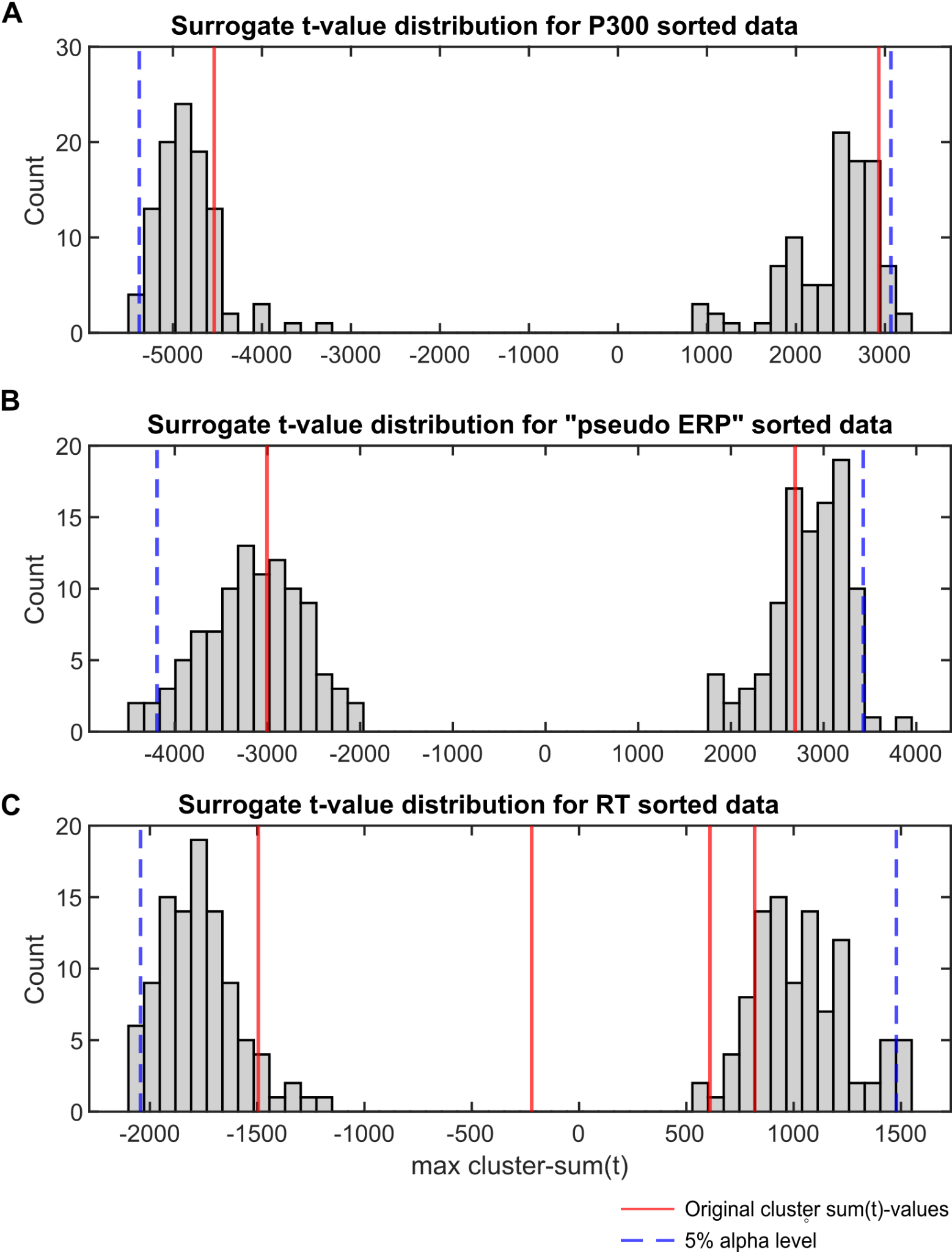

#### **Supplementary Figure 3. Non-baseline corrected data t-value distribution obtained from surrogate heartbeat control analyses.**

Histograms show summed cluster t-values for surrogate HEP differences with shuffled R-peak onsets related to trials sorted in A) high vs. low P300 trials (positive-cluster: *p* = 0.92, negative-cluster: *p* = 1), B) high vs. low “pseudo ERP” trials (positive-cluster: *p* = 0.7, negative-cluster: *p* = 0.59), and C) fast vs. slow RT trials (positive-cluster 1: *p* = 0.85, cluster 2: *p* = 1; negative-cluster 1: *p* = 0.92, *p* = 1). The red lines denote the respective cluster sum(*t*)-values of effects obtained from the original data. The dotted blue lines represent the 97.5% and 2.5% alpha cutoff. No baseline correction was performed.

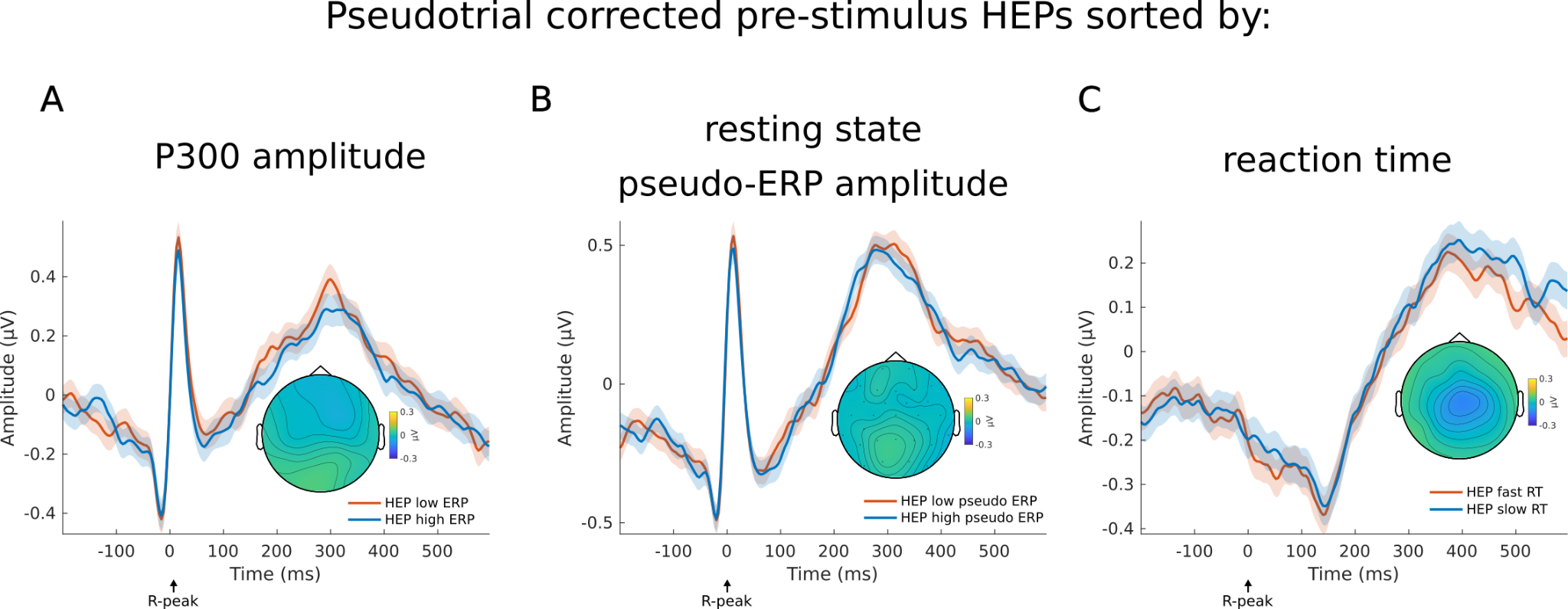

### **S3 No residual pre-stimulus HEP differences after pseudotrial correction**

After pseudotrial correction no significant condition differences for the pre-stimulus HEPs between high vs. low P300 ERP amplitude, fast vs. slow reaction times, or pseudo-ERP amplitude during resting state remain (SFig. 4). Based on the cluster of significant effects prior pseudotrial correction, HEPs were averaged and compared by equivalence testing. For all three conditions, the absence of a meaningful effect is indicated (HEPs sorted by P300: *t*_1739_ = -7.9, *p* < 0.01; “pseudo ERP” during resting state: *t*_1730_ = 7.58, *p* < 0.01; RT: *t*_1739_ = 7.18, *p* < 0.01).

#### **Supplementary Figure 4. Pre-stimulus HEPs show no meaningful differences after pseudotrial correction.**

After pseudotrial correction, no significant differences can be found for pre-stimulus HEPs that were sorted based on A) the amplitude of the subsequent P300 ERP during the oddball task , B) the amplitude of pseudo-ERPs sorted in high vs. low condition in the P300 time window during resting state, or C) the reaction times during the oddball task. The average HEPs are based on the same electrode clusters used for the respective HEPs in Figure 1 and 3. Shades around the lines are ±1 SEM.

### **S4 HEP differences between oddball task and resting state**

Before pseudotrial correction, we can only observe one cluster in positive as well as negative direction (SFig. 6A, positive cluster 1: cluster sum(*t*) = 17343, p ≤ 0.001, time: -200 – 600 ms, significant channel: Fp1, Fp2, F3, F4, F7, F8, Fz, FC1, FC2, FC5, FC6, C3, C4, T7, Cz, FT9, FT10, CP5, CP6, P3, P4, P8, Pz; negative cluster 1: cluster sum(*t*): -13498, p ≤ 0.001, time: -200 – 600 ms, significant channel: Fp1, Fp2, F4, F7, F8, T7, T8, FT9, FT10, CP5, CP6, TP9, TP10, P3, P4, P7, P8, Pz, O1, O2, PO9, PO10). Surrogate heartbeat control analysis (100 iterations) indicates that these clusters are heartbeat unrelated, as all maximum sum(*t*)-values obtained from shuffled data are larger than the original effects (positive and negative clusters *p* = 1).
Importantly, the heartbeat independent effect can be removed by pseudotrial correction, revealing several clusters distributed in time and space (the first two clusters with the largest sum(*t*) values are illustrated in SFig. 6B, for a summary of all clusters see supplementary Table 1). Surrogate heartbeat control analysis indicates that the sum(*t*)-values of all clusters fall outside of the 2.5% – 97.5% percentile range of the cluster sum(*t*)-values obtained from shuffled data (SFig. 6D). Therefore, we can conclude that the effects we observe after pseudotrial correction are coupled to the heartbeat.
To assess the potential influence of cardiac field artifacts on these differences, we compared the ECG between resting state and task by a temporal cluster-based permutation t-test. Several significant differences could be observed (SFig. 6C, p ≤ 0.001), indicating potential differences in CFA when comparing resting state and task HEPs. To further investigate if the CFA can explain the HEP differences, we tested whether the difference between resting state and task ECG ($\Delta ECG)$ correlates to the difference between resting state and task EEG ($\Delta EEG)$. No significant clusters were found in the HEP time window of interest (SFig. 7).

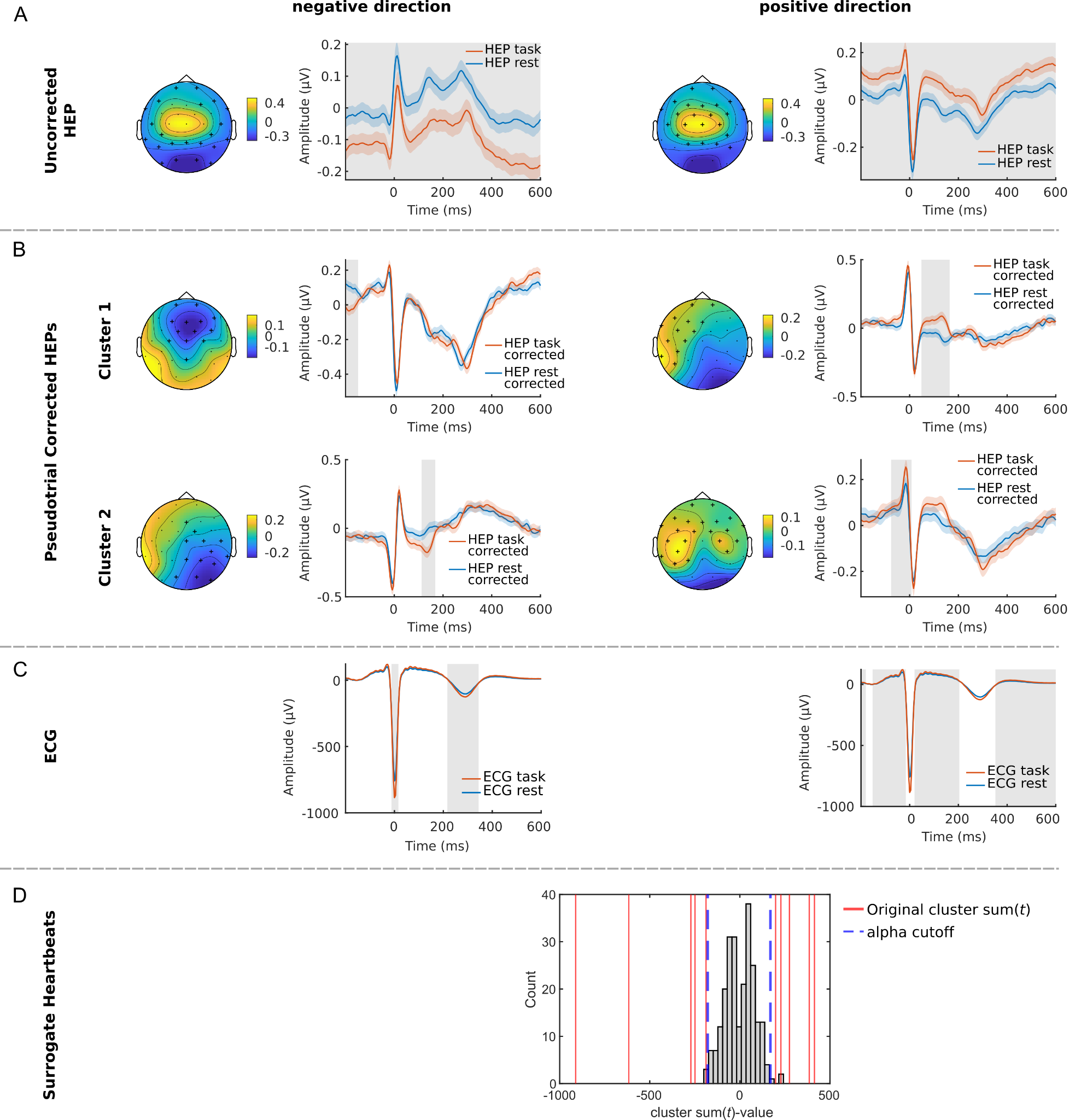

#### **Supplementary Figure 6.** Differences in HEPs between Oddball task and resting state in positive and negative direction A) before and B) after pseudotrial correction. After correction, several clusters were found and the strongest first two are illustrated. C) ECG differences between task and rest are illustrated. Shades around the lines are ±SEM, grey boxes: *p* ≤ 0.05, Topoplots are averaged over time of significant differences with crosses illustrating significant differences. D) Histogram visualizing the surrogate heartbeat control analysis. For positive as well as negative differences, all clusters of the pseudotrial corrected data have larger or smaller summed *t*-values (red lines) than the 97.5% or 2.5% alpha cutoff (blue dashed line) obtained from the largest surrogate heartbeat clusters *t*-values (gray bars).

#### **Supplementary Table 1. Significant differences between resting state and. task related HEPs after pseudotrial correction**

| **Direction** | ***p*-value** | **Summed *t*-value** | **Time Range (ms)** | **Electrodes in Cluster** |
| --- | --- | --- | --- | --- |
| positive | <= 0.001 | 782.82 | [48,164] | Fp1, Fp2, F3, F7, Fz, FC1, FC5, C3, T7, FT9, CP5, TP9, P7 |
| positive | <= 0.001 | 503.37 | [-76,8] | Fp1, Fp2, F3, F4, F7, F8, Fz, FC1, FC2, FC5, FC6, C3, C4, T7, T8, FT9, FT1,CP5, CP6, TP9, P3, P7 |
| positive | <= 0.001 | 405.94 | [532,600] | Fp1, Fp2, F3, F4, F8, Fz, FC1, FC2, FC5, FC6, C4, T8, Cz, FT1,TP10 |
| positive | <= 0.001 | 319.04 | [-200,-156] | T7, FT9, CP5, TP9, TP1,P3, P7, P8, O1, O2, PO9, PO10 |
| negative | <= 0.001 | -486 | [-200,-148] | Fp1, Fp2, F3, F4, F8, Fz, FC1, FC2, FC6, C3, C4, Cz, Pz |
| negative | <= 0.001 | -430.41 | [112,168] | Fz, FC2, C4, T8, Cz, CP6, TP1,P4, P8, Pz, O1, O2, PO10 |
| negative | <= 0.001 | -394.48 | [180,280] | F3, Fz, FC1, FC2, C3, C4, T8, Cz, CP5, CP6, TP1,P3, P4, P8, O2, PO10 |
| negative | 0.008 | -228.2 | [560,600] | C3, T7, CP5, CP6, TP9, P3, P4, P7, Pz, O1 |
| negative | 0.01 | -219.99 | [-24,20] | Fp1, F7, FC5, T7, FT9, TP9, P4, P7, P8, Pz, O1, O2, PO9, PO10 |
| negative | 0.023 | -185.04 | [292,324] | F3, F4, Fz, FC1, FC2, FC6, C4, Cz |

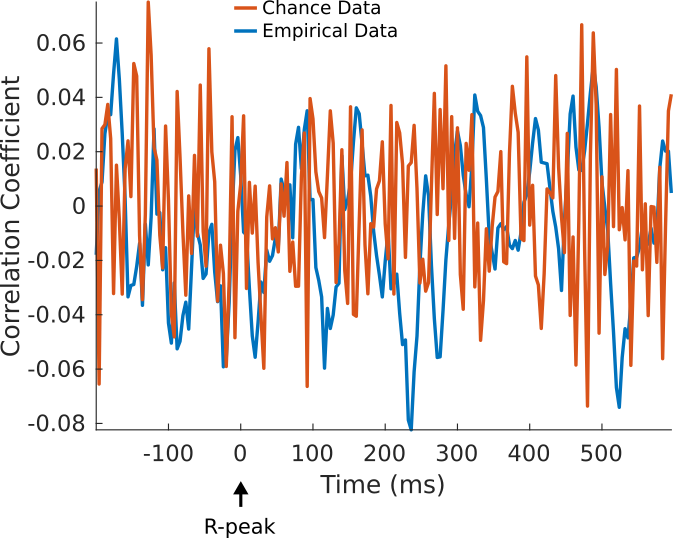

#### **Supplementary Figure 7. No Significant Correlation between** $\Delta ECG$ **and** $\Delta EEG$ **in time range of HEP analysis** Correlation between resting-state and task-related differences in ECG ($\Delta ECG$**)** and EEG ($\Delta EEG$**)** across all channels and time points (Empirical Data) and corresponding chance data obtained by correlation of $\Delta EEG$ with randomized $\Delta ECG$ data. No clusters of significant differences were found (*p*>0.05). Time courses for illustration are obtained from the Cz electrode and grand averaged across subjects. No baseline correction was performed.

**Supplementary Table 2. Clusters of significant differences between resting state and task HEPs after pseudotrial correction**

| **Direction** | **sum(*t*)-value** | ***p*-value** | **Channels** | **Time Range (ms)** |
| --- | --- | --- | --- | --- |
| positive | 414.13 | ≤ 0.001 | Fp1, Fp2, F3, F4, F7, F8, Fz, FC2, FC6, Cz, FT10 | [476, 556] |
| positive | 385.14 | ≤ 0.001 | Fp1, Fp2, F3, F4, F7, F8, Fz, FC1, FC2, FC5, T7, FT9, TP9 | [104, 160] |
| positive | 274.91 | 0.007 | Fp1, Fp2, F3, F4, F7, F8, Fz, FC1, FC2, FC5, FC6, C3, C4, T7, T8, FT10 | [-24, 12] |
| positive | 227.5 | 0.017 | Fp1, Fp2, F3, F4, F8, Fz, FC2, FC6, Cz, FT10 | [568, 600] |
| negative | -911.71 | ≤ 0.001 | F3, FC1, FC5, C3, T7, FT9, CP5, CP6, TP9, P3, P4, P7, P8, Pz | [368, 600] |
| negative | -616.67 | ≤ 0.001 | FC1, C3, C4, T8, Cz, CP5, CP6, TP10, P3, P4, P7, P8, Pz, O1, O2, PO10 | [112, 240] |
| negative | -271.89 | 0.003 | F3, F4, Fz, FC1, FC2, FC6, C3, C4, Cz | [-200, -160] |
| negative | -249.27 | 0.004 | F7, FC5, C3, T7, FT9, CP5, TP9, P3, P7, P8, O1, O2, PO9, PO10 | [-20, 24] |
| negative | -187.81 | 0.023 | F3, FC1, C3, CP5, TP9, P7 | [260, 332] |
